# Novel aspects of iron homeostasis in pathogenic bloodstream form *Trypanosoma brucei*

**DOI:** 10.1101/2021.05.27.445749

**Authors:** Carla Gilabert Carbajo, Lucy J. Cornell, Youssef Madbouly, Zhihao Lai, Phillip A. Yates, Michele Tinti, Calvin Tiengwe

## Abstract

Iron is an essential regulatory signal for virulence factors in many pathogens. Mammals and bloodstream form (BSF) *Trypanosoma brucei* obtain iron by receptor-mediated endocytosis of transferrin bound to receptors (TfR) but the mechanisms by which *T. brucei* subsequently handles iron remains enigmatic. Here, we analyse the transcriptome of *T. brucei* cultured in iron-rich and iron-poor conditions. We show that adaptation to iron-deprivation induces upregulation of TfR, a cohort of parasite-specific genes (ESAG3, PAGS), genes involved in glucose uptake and glycolysis (THT1 and hexokinase), endocytosis (Phosphatidic Acid Phosphatase, PAP2), and most notably a divergent RNA binding protein RBP5, indicative of a non-canonical mechanism for regulating intracellular iron levels. We show that cells depleted of TfR by RNA silencing import free iron as a compensatory survival strategy. The TfR and RBP5 iron response are reversible by genetic complementation, the response kinetics are similar, but the regulatory mechanisms are distinct. Increased TfR protein is due to increased mRNA. Increased RBP5 expression, however, occurs by a post-transcriptional feedback mechanism whereby RBP5 interacts with its own, and with *PAP2* mRNAs. Further observations suggest that increased RBP5 expression in iron-deprived cells has a maximum threshold as ectopic overexpression above this threshold disrupts normal cell cycle progression resulting in an accumulation of anucleate cells and cells in G2/M phase. This phenotype is not observed with overexpression of RPB5 containing a point mutation (F61A) in its single RNA Recognition Motif. Our experiments shed new light on how *T. brucei* BSFs reorganise their transcriptome to deal with iron stress revealing the first iron responsive RNA binding protein that is co-regulated with TfR, is important for cell viability and iron homeostasis; two essential processes for successful proliferation.

**Author Summary:** African trypanosomes are single-celled extracellular parasites of humans and animals relying on essential host nutrients for survival. They satisfy their iron needs by capturing host transferrin-bound iron using a surface-localised transferrin receptor (TfR) that is structurally distinct from its host counterpart. Little is known about the trypanosome response to fluctuations in host iron availability, with the exception of modulated TfR expression. We show that unlike other eukaryotes, at the transcriptome level, trypanosomes do not regulate iron-dependent enzymes as a mechanism to cope with iron deprivation.

Instead, we identify a group of novel iron responsive trypanosome-specific genes, particularly an RNA Binding Protein RBP5 that is responsive to iron levels, albeit mediated by a distinct mechanism from TfR. We show that although RBP5 expression is elevated at the mRNA and protein levels, increased abundance above a maximum threshold is toxic. The trypanosome TfR has been suggested as a therapeutic target, but whether it is essential for optimal host colonisation is unclear. Our data demonstrate that trypanosomes efficiently import free iron from their environment independent of TfR suggesting that alternative iron uptake pathways exist, and that any therapeutic interventions targeting TfR must be evaluated with caution.

## Introduction

One remarkable challenge for invading pathogens is the ability to compete for and sequester host nutrients. Because nutrient availability often fluctuates, pathogens co-opt diverse strategies to sense, respond and successfully adapt within their hosts. This is especially significant for African trypanosomes, single-celled Kinetoplastid parasites that colonise various niches within their mammalian hosts (1, 2) and the midgut of tsetse-fly vectors. Several species of the *Trypanosoma* genus are pathogenic to wild and domestic animals, of which two sub-species *T. brucei gambiense* and *T. b. rhodesiense* infect humans.

Bloodstream form (BSF) trypanosomes rely exclusively on host iron for survival. Elemental iron is an essential co-factor in many biological reactions, but toxicity associated with “free iron” creates the need for tight regulation (3–5). In vertebrates, extracellular ferric iron is bound by transferrin to avoid toxicity. *T. brucei* sequesters host transferrin-bound iron (Tf/Fe^3+^) from the bloodstream using a divergent high affinity transferrin receptor (TfR) (6, 7). TfR is expressed from one of ∼15 sub-telomeric bloodstream expression sites (BES) with genes important for antigenic variation (8–10). Functional TfR is heterodimeric encoded by two *E*xpression *S*ite *A*ssociated *G*enes: *ESAG6* (*E6*) and *ESAG7* (*E7*) sharing 80% amino acid sequence identity and a short variable domain critical for transferrin (Tf) binding (11) or antigenic variation (6). The heterodimer localizes to endosomes and the flagellar pocket (FP); a membrane invagination for endo- and exo-cytosis (7, 9). TfR delivers Tf/Fe^3+^ by receptor-mediated endocytosis to the lysosome where Fe^3+^ is reduced to Fe^2+^ by unknown ferric reductases and subsequently transported into the cytosol by a mucolipin-like protein (12, 13). Released Fe^2+^ is either coordinated with iron-dependent proteins during protein synthesis or stored by yet unknown machinery. The intracellular iron storage protein ferritin has not been found in trypanosomes. Tf is ultimately degraded by lysosomal Cathepsin B proteases (14), but whether TfR is recycled remains contentious (6, 15), and how it is regulated is currently unknown.

In mammalian cells, TfR regulation by iron involves post-transcriptional feedback mechanisms whereby iron-regulatory proteins (IRPs) bind iron-responsive elements (IREs) in the 3’-UTRs of mRNAs (16, 17). When iron is low, aconitase (also known as IRP1) binds IREs in the TfR 3’-UTR, stabilises its mRNA and enhances translation increasing iron uptake (18). Conversely, when in excess, iron binds iron-sulfur clusters in IRP1 releasing it from the TfR 3’-UTR, leading to degradation by exonucleases (16, 19).

Several divergent biological features of *T. brucei* suggest that iron regulation is distinct. First, insect stage procyclic forms (PCF) do not express TfR since BES are transcriptionally silenced. Iron uptake occurs by a two-step reductive process of ferric complexes (20), or by an alternative heme transporter, TbHrg (21). Second, although BSFs express a haptoglobin-hemoglobin receptor HpHbR (22), absent in PCFs, they lack an identifiable homologue of heme oxygenase required to breakdown heme into usable Fe^2+^. Imported heme is directly incorporated into proteins implying HpHbR does not provide a source for elemental iron (23). Third, BSF *T. brucei* lack most cytochromes since energy production occurs primarily by glycolysis in the glycosome (24). Nevertheless, putative orthologues of iron-dependent enzymes are found in both lifecycle stages, notably aconitase, ribonucleotide reductase, alternative terminal oxidase and superoxide dismutase, two ferric reductases and multiple iron transporters (25). Whether these genes are regulated by iron like their vertebrate hosts is unknown. Finally, trypanosome TfR is transcribed by RNA Pol I, an enzyme normally dedicated for rRNA transcription in eukaryotes, signifying an interesting departure from eukaryotic models for maintaining iron homeostasis.

Despite these distinctions, culturing BSF *T. brucei* cells in canine serum (26–28), under hypoxic conditions (27), in TfR-depleted cells (29, 30), or treatment with the iron chelator Deferoxamine (DFO) (31) result in 3 –10-fold upregulation of TfR mRNA and protein, suggesting that they respond to environmental fluctuations in iron levels. In the latter scenario, gene deletion of aconitase did not affect iron-induced TfR upregulation, demonstrating that TfR regulation occurs independently of aconitase, and that *T. brucei* lacks a canonical IRP/IRE-mediated regulatory system (31).

To investigate outstanding questions on iron-dependent control of TfR and study how mammalian-infective trypanosomes adapt to iron deficiency, we employed two approaches: (i) short term treatment with DFO to mimic depletion of the cellular iron pool and (ii) RNA silencing of TfR to block Tf uptake. We discover a cohort of specific iron responsive genes including a novel Kinetoplastid-specific RNA Binding Protein RBP5 that is co-regulated with TfR, but by a distinct mechanism. We define the precise level of iron-dependent post-transcriptional control of TfR and RBP5 expression, determine the kinetics of the iron response and show that exogenous iron can compensate for TfR loss. To our knowledge, this is the first report of a conserved Kinetoplastid RBP that is responsive to iron levels.

## Results

### TfR is regulated in response to intracellular iron status

To establish the specificity of TfR regulation in iron-limiting and iron-replete conditions, we incubated wild type BSF cells with increasing concentrations of deferoxamine DFO (0 - 25 μM), Iron (III) chloride (FeCl_3_, 0 - 25 μM) or with sub-lethal doses of iron-presaturated deferoxamine (DF, 10:12.5 μM) over a 24-hr period as specificity control (**Fig 1a**). The drugs and doses had either been used by previous investigators (32–34) or were empirically determined. In our hands, DFO inhibited growth by 25%, 50% and 100% at 24 hr of incubation with 5, 10, and 25 μM concentrations, respectively (**left**). Ferric iron was not toxic at lower concentrations but caused a slight growth inhibition at 25 μM over a 24-hr period (**middle**). No significant growth defect was observed with 2.5 μM excess iron pre-saturated with DFO (**right**). Our growth measurements under these conditions indicate that iron limitation by DFO prevents normal cell proliferation and exogenous Fe (III) at the chosen doses does not significantly affect cell viability.

**Figure 1.**
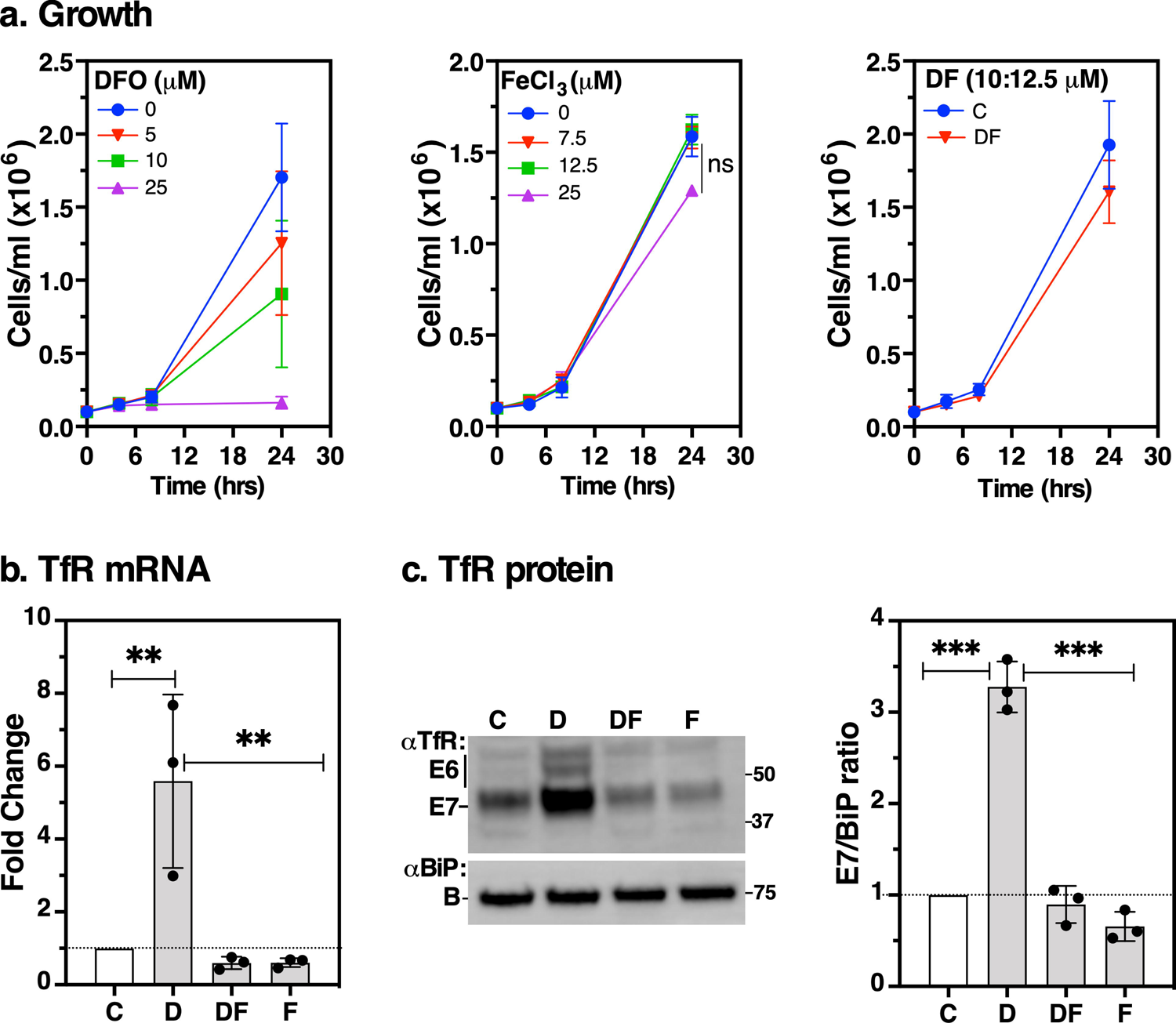
Effect of deferoxamine or ferric iron on cell viability and TfR expression. Log phase BSF cells (∼5×10^5^ cells/ml) were incubated with the iron chelator deferoxamine (DFO, D), iron-saturated deferoxamine (DF), Iron (III) chloride (FeCl_3_, F) or untreated (C). **a.** Cell densities (mean ± SD, *n* = 3) were determined by haemocytometer in the presence of varying concentrations of DFO, FeCl_3_ or DF as shown in the insets; ns: not significant. **b.** RNA was harvested from cells following 5 hr treatments with D (25 μM); DF (25:27.5 μM); F (25 μM) or untreated (C). *TfR* mRNA levels were determined by qRT-PCR normalised to *ZFP3* signal and expressed as a ratio to the control treatment values. Data are means ± SD (*n* = 3 biological replicates; with 3 technical replicates for each *n*). *\*\*P* < 0.002, as determined by one-way ANOVA with post-hoc Tukey test. **c.** Protein levels were detected by immunoblotting with anti-TfR (αTfR) or anti-BiP (αBiP) as loading control. Bar chart (right) shows signals of E7 from αTfR immunoblot quantified by densitometry using Image Lab software (Bio-Rad) normalised to the corresponding BiP signals. Data are means ± SD (*n* = 3 biological replicates). \**\*\*P* < 0.001, as determined by one-way ANOVA with post-hoc Tukey test.

Next, we re-evaluated the effect of iron on *TfR* mRNA and protein levels by acute treatment with the highest doses of each of these compounds (either alone or in combination) at time intervals compatible with cell viability. Treatment with DFO (25 μM, 5 hr) alone resulted in ∼5-fold and 3-fold increase in endogenous *TfR* mRNA and protein levels, respectively (**Fig 1b** and **1c**). Conversely, compared to untreated controls, TfR transcript and protein decreased by ∼50% and ∼30% when cells were treated with excess FeCl_3_. We conclude *TfR* mRNA and protein are upregulated in iron-limiting and downregulated in iron-saturating conditions.

### Iron starvation results in increased *TfR* mRNA stability and enhanced translation

To determine the level of iron-dependent post-transcriptional control of TfR, we incubated wild type cells with DFO followed by addition of sinefungin to block trans-splicing and thus mRNA maturation, and actinomycin D to inhibit transcription. We performed qRT-PCR at various time points over a 2-hr period and found that *TfR* (*E7, E6*) message levels decayed more slowly in DFO-treated compared to control cells (**Fig 2a**). The half-lives of *E7* and *E6* were 24.8 and 25.8 min, respectively, in control cells but increased to 65.4 and 73.1 min, respectively, in DFO-treated cells. *Actin* mRNA half-life measurements from the same cDNA samples were 20 and 15.7 min in DFO-treated and control cells, respectively (**Fig 2b**). The difference in *Actin* t_1/2_ was not statistically significant and matched earlier reports (35), validating our TfR half-life measurements.

**Figure 2.**
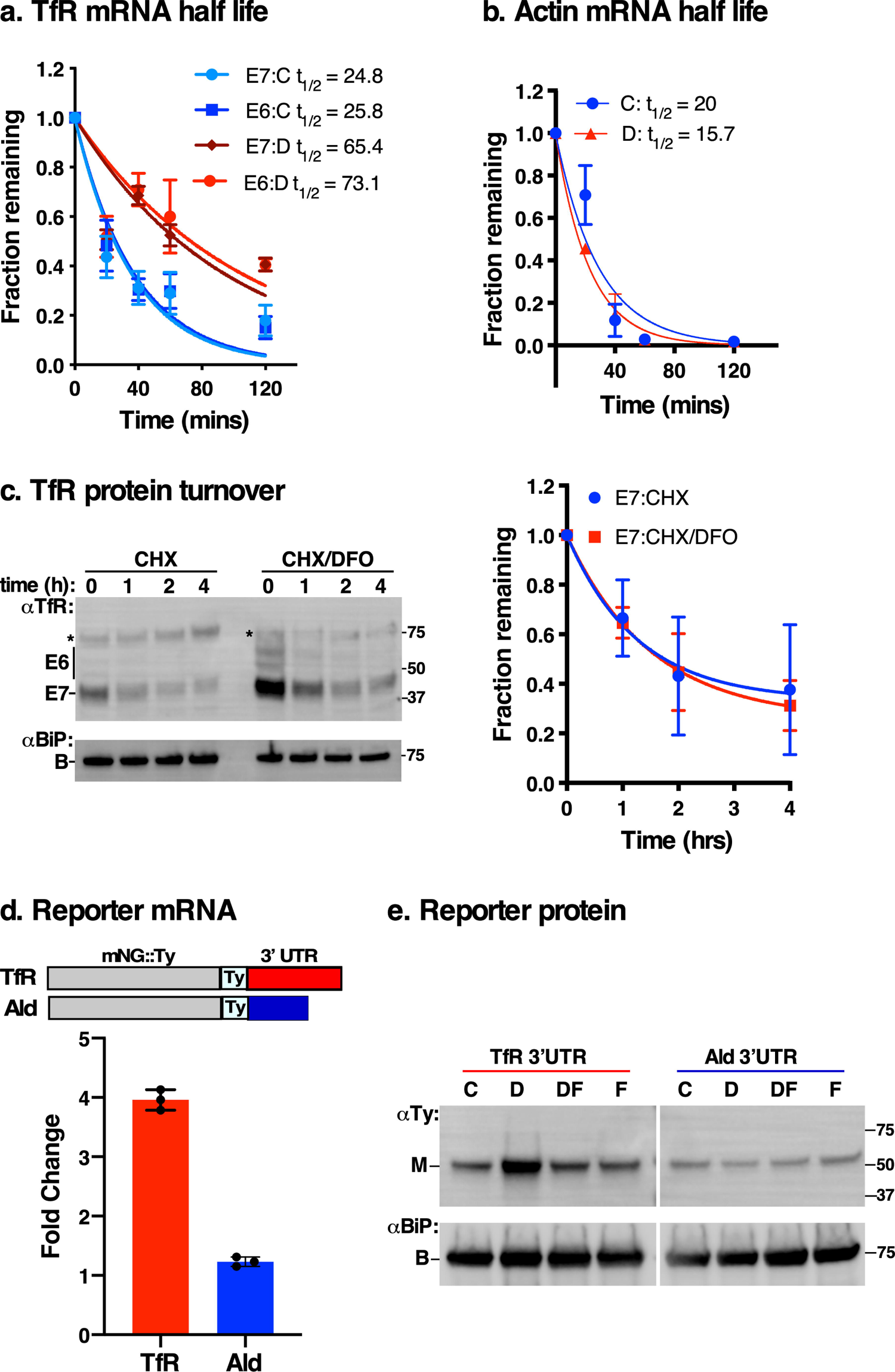
TfR turnover kinetics following iron starvation. **a.** Log phase cells (∼5×10^5^ cells/ml) were pre-treated with DFO (D) or untreated (C) for 5 hr and then incubated with sinefungin (5 μg/ml, 5 min) to block trans-splicing and actinomycin D (10 μg/ml) to inhibit transcription. Total RNA was isolated at specified time intervals and relative mRNA levels of *E6* & *E7* were determined by qRT-PCR normalised to 28S ribosomal RNA as internal control. Values plotted on graphs are normalised to that of time zero. The estimated half-life (t_1/2_, min) was calculated based on regression analysis using PRISM. Data are means ± SD (*n* = 3 biological replicates; with 3 technical replicates for each *n*). **b.** The half-life of *Actin* was determined using cDNA samples in (**a**). Actin serves as a control for t_1/2_ measurements. Data are means ± SD (*n* = 3 biological replicates; with 3 technical replicates for each *n*). **c.** Cells were incubated with 100 μg/ml cycloheximide (CHX) to block protein synthesis following pre-treatment with DFO (CHX/DFO, 5 hr) or without (CHX). Protein levels were analysed at time intervals indicated by immunoblotting with αTfR or αBiP (loading control). Star indicates a non-specific background polypeptide. Decay curve (right) shows rate of E7 protein turnover quantified from immunoblots with CHX/DFO treatment (E7:CHX/DFO) or CHX only (E7:CHX), normalised to the corresponding BiP signals at indicated time points relative to that of time zero. Data are means ± SD (*n* = 3 biological replicates). **d.** *TfR 3’– UTR* confers stability to mNeonGreen reporter in iron starvation conditions. Cells expressing mNeonGreen::Ty (mNG::Ty) reporter fused to either *E6* (*TfR*, red) or *Aldolase* (blue) 3’-UTR were treated with Deferoxamine (D) for 5 hr, total RNA was extracted and relative mNG transcript levels quantified by qRT-PCR, normalised to untreated controls with *ZFP3* as endogenous standard. Data are means ± SD (*n* = 3 technical replicates, with 3 technical replicates for each *n*). **e.** Western blot analysis of mNG::Ty protein with anti-Ty (αTy) or anti-BiP (αBiP) antibodies (loading control) from whole cell extracts treated with Deferoxamine (D), iron saturated deferoxamine (DF), FeCl_3_ (F) or left untreated (C), as previously described. Blot is representative of *n* = 3 independent biological replicates.

Previous studies on TfR regulation overlooked the potential contribution of post translational mechanisms. We investigated this issue by pre-treating trypanosomes with DFO to induce TfR upregulation, supplemented the cultures with 100 μg/ml cycloheximide (CHX) to block protein synthesis, and then quantified TfR protein levels over time from anti-TfR immunoblots (**Fig 2c**). While TfR protein increased following DFO treatment relative to controls (compare t = 0, CHX vs CHX/DFO), the turnover rate remained unchanged, indicating that increased translation due to increased mRNA stability drives upregulation in low iron conditions. Finally, we tethered the *TfR 3’-UTR* to an mNeonGreen (mNG) reporter construct (**Figs 2d & 2e**). Following DFO treatment, the behaviour of mNG reporter mRNA and protein mimicked endogenous *TfR* in **Fig. 1**. These results argue that iron-dependent stability of *TfR* mRNA relies on putative conserved *cis*-acting IREs in its 3’-UTR, as previously reported (33); confirming its role in iron-dependent regulation.

### Transcriptomics identifies iron regulated genes in BSF *T. brucei*

To identify iron regulated genes, we treated wild type cells with DFO (D) or FeCl_3_ (F) either alone or in combination (DF), performed mRNA-Seq in triplicates per condition, and compared them to controls (C). The analysis is provided in an online visualisation tool at https://calvin-tfr.onrender.com/FiguresShare.html. Applying a threshold of ≥2-fold (FDR < 0.05), *E6*/*E7* mRNAs were ∼4-fold upregulated in all DFO-treated conditions (**Fig 3a**), consistent with qRT-PCR data (**Fig 1b**) and other reports (30, 31, 33); validating the transcriptomic analyses. We found no significant change in expression of iron-dependent enzymes, a phenomenon typically observed in other systems (36), suggesting that trypanosomes have co-opted alternative mechanisms for regulating iron. Apart from *TfR*, the most significantly upregulated gene was an uncharacterised putative RNA Binding protein RBP5. The Venn diagram shows the distribution of the number of upregulated transcripts among all the treatment conditions (**Fig 3b**). All differentially regulated transcripts are in *Supplementary Tables S1*.

**Figure 3.**
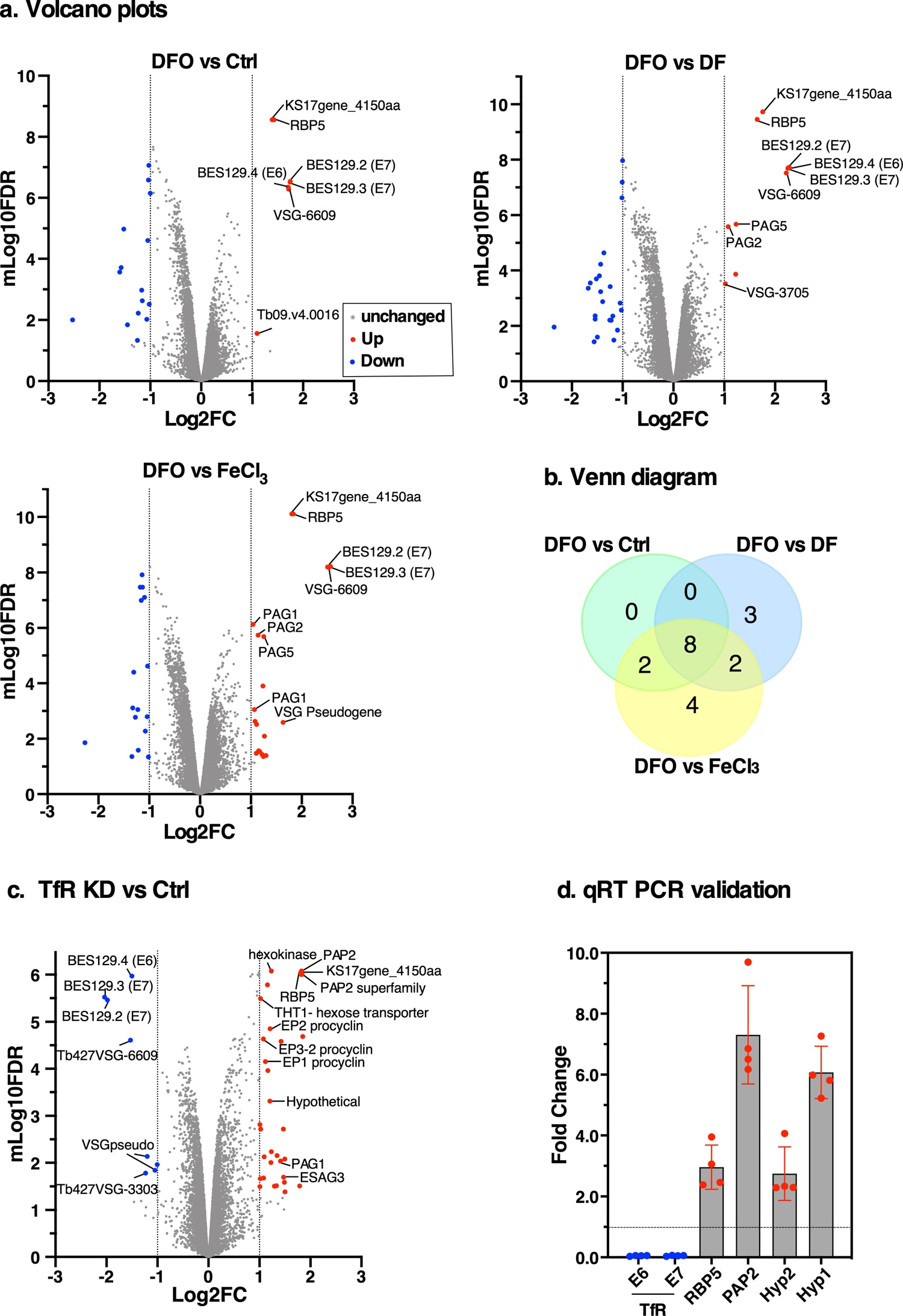
Transcriptional profiling of iron regulated genes in BSF *T. brucei*. **a.** Effect of DFO on the transcriptome. Log phase cells (∼5×10^5^ cells/ml) were incubated with 25 μM DFO, 25 μM DFO pre-saturated with 27.5 μM FeCl_3_ (DF), 25 μM FeCl_3_ or untreated (Ctrl). Total RNA was purified from three technical replicates for each condition and subjected to RNA sequencing. Volcano plots show transcriptome data of differentially expressed genes with 2-fold change relative to controls (FDR adjusted p-value < 0.05): grey dots (no change), red dots (upregulated), blue dots (downregulated). All differentially expressed gene IDs are presented as *supplementary Table S1*. **b.** The Venn diagram shows the distribution of the number of differentially expressed genes in each condition shown in (a) above. **c.** Transcriptome analysis of cells following iron starvation by TfR silencing. Total RNA was purified from TfR knockdown (TfR KD) or control (Ctrl) cells and RNA sequencing was performed. Volcano plot showing transcriptome data (*n* = 3 technical replicates) following differential expression analyses: grey dots (no change), red dots (upregulated), blue dots (downregulated). Differentially expressed genes with (FDR adjusted p-value < 0.05) are coloured as in (2a). >2-fold change relative to controls **d.** qRT-PCR was performed to independently validate selected upregulated transcripts from the RNA-Seq in (**c**). Relative mRNA levels of endogenous TfR subunits *ESAG6 (E6), ESAG7(E7)*, RNA Binding Protein 5 (*RBP5*), Phosphatidic Acid Phosphatase (*PAP2*), and mRNAs of two hypothetical proteins (Hyp2, Tb927.8.510 and Hyp1, Tb927.8.490) are shown. mRNA levels are normalised to the corresponding *ZFP3* signal and expressed as a ratio to the control treatment values. Data are means ± SD (*n* = 4 biological replicates; with 3 technical replicates for each *n*).

Next, we investigated the effects of iron starvation by tetracycline inducible RNA silencing of TfR. Previously, we showed that TfR silencing impaired iron uptake (30). Here, we recreated the TfR RNAi cells and replicated the previous results (**Fig S1, a - d**). We then induced RNAi (24 hr), quantified mRNA levels of selected homologues of iron-dependent genes in *T. brucei* (**Fig S1e**) and found no difference in their steady state levels as judged by qRT-PCR, consistent with data in **Fig 3a**. To identify other differentially regulated genes, we deep-sequenced mRNA from three technical replicates and show the analyses at https://calvin-tfr2.onrender.com/FiguresPaper.html. Forty genes showed >2-fold (FDR < 0.05) regulation – **Fig 3c** and *Supplementary Table S1*). Among these were RNA-binding protein (RBP5), five hypothetical proteins, two putative phosphatidic acid phosphatase related genes (PAP2), seven VSG-related genes from silent Bloodstream Expression Sites (BES), hexokinase and a hexose transporter (THT1), ESAG3, three Procyclin genes (EP1-3), and two Procyclin-Associated Genes (PAGS). Apart from three *VSG*-related genes and one “novel” transcript, all down-regulated genes were subunits of TfR, validating our data.

Next, we quantified four of the most significantly upregulated genes by qRT-PCR following TfR knockdown and found a ∼90% reduction in *E6/E7* transcripts, upregulation of *RBP5* (∼3-fold), *PAP2* (∼7-fold) and two hypothetical genes (∼3-fold and ∼6-fold), respectively (**Fig 3d**). Collectively, these results suggest that blocking iron uptake by TfR silencing has no direct effect on mRNA expression levels of iron-dependent enzymes but results in modulation of exclusively trypanosome-specific genes (VSGs, ESAG3, RBP5, PAGS, EP1-3), genes involved in glycolysis (hexokinase and THT1), and endocytosis (PAP2) supporting a divergent mechanism for regulating iron.

### Iron supplementation rescues growth and abolishes the iron starvation response

Our experiments showing that free FeCl_3_ modulates TfR expression suggest that trypanosomes can acquire and utilise exogenous iron. To test if exogenous iron entered the cells through the TfR/Tf complex or by an alternative route, and if this was sufficient to rescue the growth defect of TfR-depleted cells, we adopted an RNAi-based knock-in system in which an RNAi-resistant TfR subunit *E6^R^* was expressed in the parental RNAi background (30). Briefly, the *E6^R^* reporter construct replaced the CDS of native *E6* but preserved endogenous intergenic regions which would be expected to mediate an iron starvation response when native TfR is silenced. **Fig 4a** shows that RNAi induction with tetracycline resulted in ∼90% reduction in *E6/E7* native transcripts and 3-fold increase in *E6^R^* mRNA as predicted (30). Cell growth was then analysed in the presence of tetracycline supplemented with 200 μg/ml holo-Tf, 200 μg/ml lactoferrin, or 25 μM FeCl_3_ (**Fig 4b**). Tetracycline alone inhibited growth and addition of either holo-Tf or lactoferrin had no effect, consistent with a lack of functional TfR or compensation by a lactoferrin binding protein, respectively.

**Figure 4.**
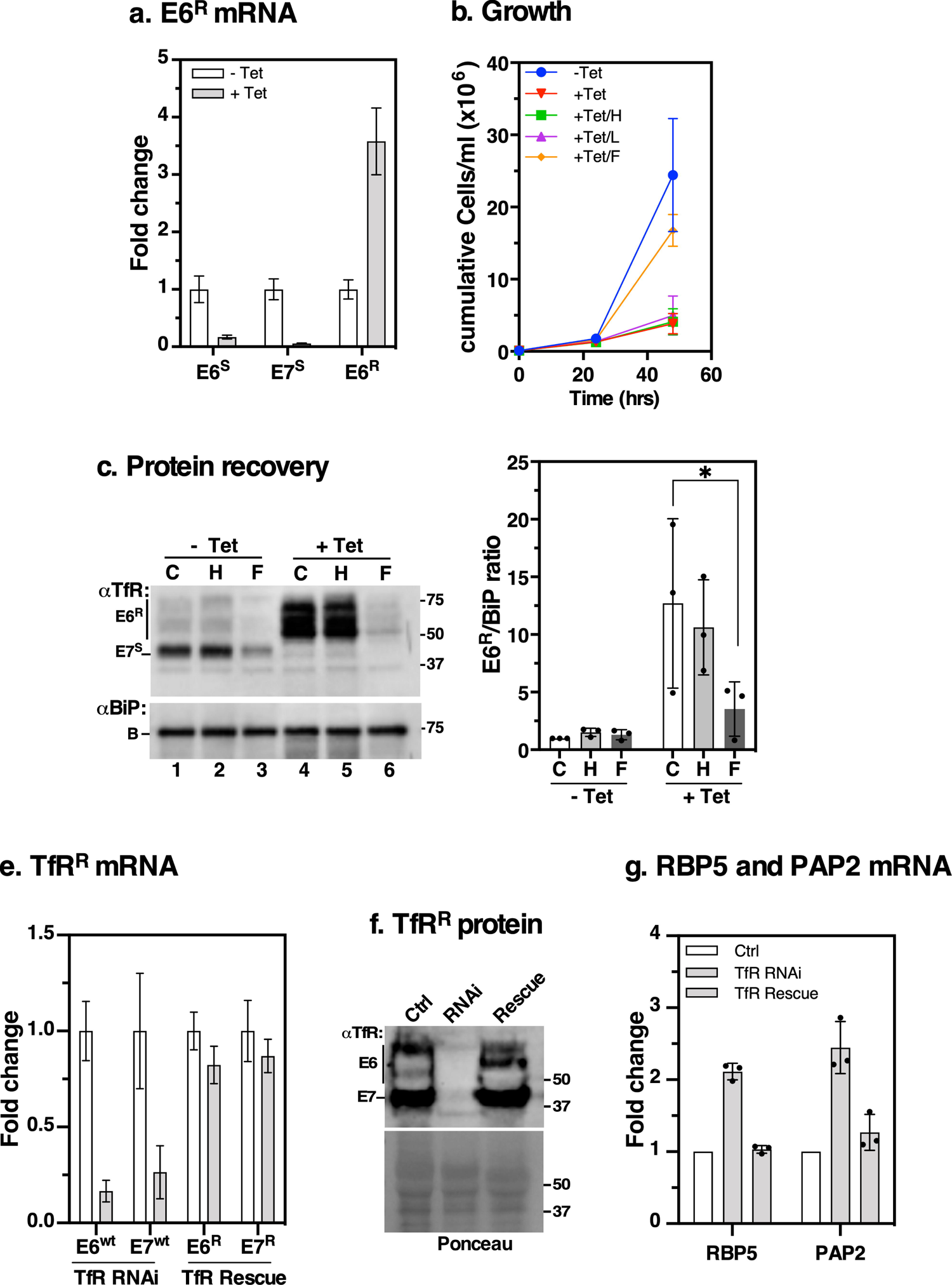
Effect of exogenous iron on growth and E6^R^ expression in TfR-depleted cells. **a.** Generation of RNAi-resistant E6^R^ reporter cell line to monitor the iron starvation response following TfR silencing. Parental TfR RNAi cells were transfected with an RNAi-resistant E6 (*E6^R^*) construct replacing native E6 CDS but retaining its native 5’- and 3’-UTRs. RNAi was induced with tetracycline (+Tet, 24 hr), and mRNA levels of RNAi-sensitive *E6^S^, E7^S^*, and RNAi-resistant *E6^R^* were determined by qRT-PCR. *E6^S^* transcripts derive from silent BESs while *E7^S^* derive from both active and silent BESs. Fold change is calculated relative to respective un-induced controls (-Tet) with *ZFP3* as endogenous control. **b.** Cumulative growth of E6^R^ expressing cells cultured in the presence (+ Tet) of 1 μg/ml tetracycline supplemented with holo-transferrin (H, 200 μg/ml), Lactoferrin (L, 200 μg/ml), FeCl_3_ (F, 25 μM) or untreated (C). Counts are means ± SD (*n* = 3 biological replicates). **c.** 10^7^ cell equivalents/condition from (b) were harvested 24 hr after treatments, total protein was blotted with αTfR or αBiP; excluding lactoferrin samples. Bar chart (right) shows signals of E6^R^ quantified by densitometry using Image Lab software (BioRad). E6^R^ signals/condition is normalised to the corresponding BiP signal and expressed as a ratio relative to un-induced control (-Tet/C). Western blot is representative of *n* = 3 biological replicates and plot shows means ± SD. *\*P* < 0.05, as determined by two-way ANOVA with post-hoc Tukey test. **e, f**. The E6^R^ reporter cell line was transfected with RNAi-resistant *E7^R^* to reconstitute functional RNAi-resistant TfR in parental RNAi background. RNAi was induced in parental RNAi-sensitive (TfR RNAi) or RNAi-resistant TfR (TfR Rescue) cells. Relative levels of *E6/E7* mRNA (**e**) or protein (**f**) were analysed by qRT-PCR or by immunoblotting with αTfR, respectively. Ponceau stained membrane served as loading control. **g.** Expression of *RBP5* and *PAP2* with or without functional complementation of TfR. Relative mRNA levels of *RBP5* and *PAP2* were analysed by qRT-PCR following TfR silencing in RNAi-sensitive cells without (TfR RNAi) or with (TfR Rescue) functional RNAi-resistant versions. All signals are normalised to un-induced cells relative to *ZFP3* as endogenous control. Data are means ± SD (*n* = 3).

Surprisingly, supplementation with exogenous FeCl_3_ partially rescued growth. Using protein extracts derived from the treated cells in **Fig. 4b** (excluding lactoferrin samples), anti-TfR western blots showed ∼10-fold upregulation in E6^R^ reporter expression (**Fig 4c**, compare lanes 1 & 4; 2 & 5) in Tet/control and Tet/holo-Tf treatments, which decreased by ∼70% during incubation with FeCl_3_ (compare lanes 4 & 5 vs lane 6). Collectively, these data indicate that BSF *T. brucei* acquired exogenous free iron independent of TfR and this was sufficient to rescue growth and terminate iron starvation response signals. Similar results were obtained with 48 hr treatments, or when the cells were pretreated with tetracycline for 24 hr followed by supplementation with FeCl_3_ (**Fig S2**).

### Genetic complementation restores normal expression of *RBP5* and *PAP2*

Next, we validated the specificity of the iron starvation response by transfecting RNAi-resistant *E7^R^* constructs into the E6^R^ expressing strain to functionally complement TfR in the RNAi cells. Using the parental RNAi and reconstituted strains, treatment with tetracycline for 24 hr caused a ∼80% decrease in native *E6/E7* transcripts (**Fig 4e**, TfR RNAi) which was restored to wild type levels by constitutive expression of resistant E6^R^/E7^R^ (**Fig 4e**, TfR Rescue). The same pattern was seen for protein expression as judged by anti-TfR western blotting (**Fig 4f**). In **Fig 4g**, we quantified *RBP5* and *PAP2* mRNA levels in TfR RNAi cells with or without reconstituted RNAi-resistant TfR. We found ∼2-fold upregulation of *RBP5* and *PAP2* in RNAi cells which was restored to wild-type levels in the RNAi-resistant cells. These data confirm that the RNAi-resistant versions could complement the iron transport functions of native TfR and abrogate the iron response of *RBP5* and *PAP2* mRNAs.

### RBP5 is a Kinetoplastid-specific protein that localises to the cytosol

Next, we focused our attention on RBP5 (Tb927.11.12100) since it was upregulated along with TfR in both iron-depleted conditions. RBP5 has a canonical RNA Recognition Motif (RRM) between residues 16 and 86 as predicted by PFAM (**Fig 5a**). Phylogenetic analyses showed that it is a conserved Kinetoplastid-specific protein with no apparent eukaryotic homologues (**Fig 5a & S3**), indicative of an evolutionarily conserved function.

**Figure 5.**
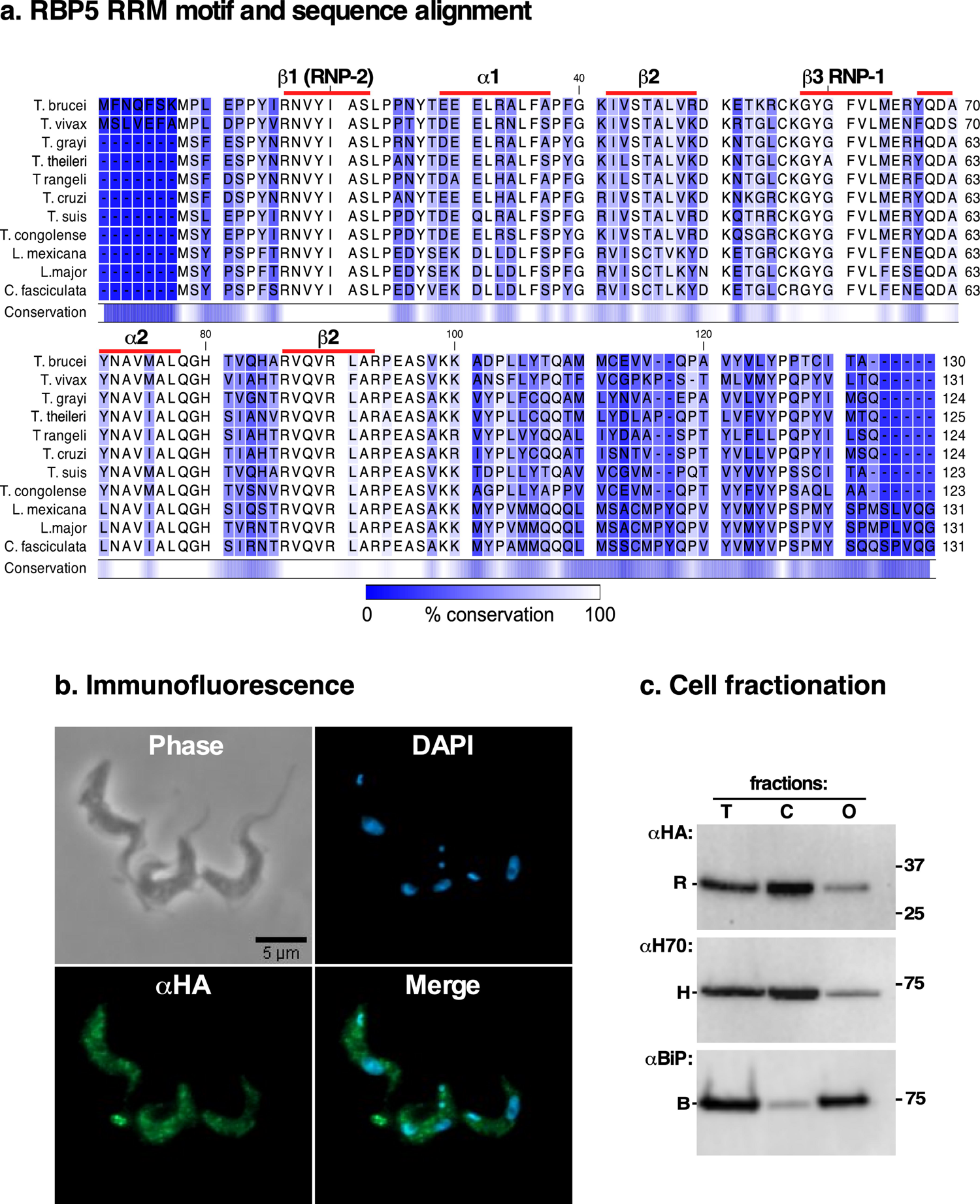
RBP5 domain architecture, sequence alignment and localisation. **a.** RBP5 sequences from *Trypanosoma brucei, T. vivax, T. grayi, T. rangeli, T. congolense, T. theileri, T. cruzi, T. suis, Leishmania major, L. mexicana,* and *Crithidia fasciculata* were aligned using CLC Genomics Workbench (Qiagen). The blue shading highlights percent conservation according to the bar below. Secondary structure elements depicting its single RNA Recognition Motif (RRM) are highlighted by red lines above respective amino acid residues as predicted by PFAM; (e-value 1.6e-18). **b.** Immunofluorescence staining of HA::RBP5 epitope-tagged cells visualized with αHA (green; RBP5) and DAPI (blue; nucleus and kinetoplast DNA). **c.** HA::RBP5 cells were hypotonically lysed and fractionated by centrifugation into total (T), cytosolic (C) or organelle associated (O) fractions, separated by SDS-PAGE and immunoblotted with respective antibodies for HA::RBP5 (αHA, R), endogenous BiP (αBiP, B, membrane) and Hsp70 (αH70, H, cytosol), respectively. Note: residual contamination of Hsp70 in organelle-associated fraction; most likely due to incomplete cell membrane disruption.

Immunofluorescence (IF) microscopy on fixed, permeabilised HA::RBP5 epitope-tagged cells with anti-HA antibodies revealed that it was cytosolic (**Fig 5b**) with no obvious nuclear localisation. This observation was confirmed by cell fractionation experiments using the ER molecular chaperone BiP and Hsp70 as markers of organelle-associated and cytosolic proteins respectively (**Fig 5c**).

### RBP5 is regulated by cellular iron levels and is unstable in iron replete conditions

To verify RBP5 regulation by iron we performed qRT-PCR analysis in wild type cells. Endogenous *RBP5* mRNA increased ∼3-fold during DFO incubation and slightly decreased (∼30%) with either iron presaturated DFO or with FeCl_3_ relative to untreated controls (**Fig 6a**). To assess if a similar effect occurred at the protein level, we fused both alleles of RBP5 *in situ* with a 6xHA tag on the N-terminus using CRISPR/Cas9 (37) ensuring constitutive expression with its native 3’-UTR (**Fig S4a**). Western blot analyses of HA::RBP5 expression following DFO or FeCl_3_ treatment showed ∼8-fold upregulation and 90% downregulation, respectively (**Fig 6b**), confirming that both mRNA and protein are regulated by iron. Since both RBP5 alleles were epitope tagged, it demonstrated that epitope tagging did not interfere with the iron response.

**Figure 6.**
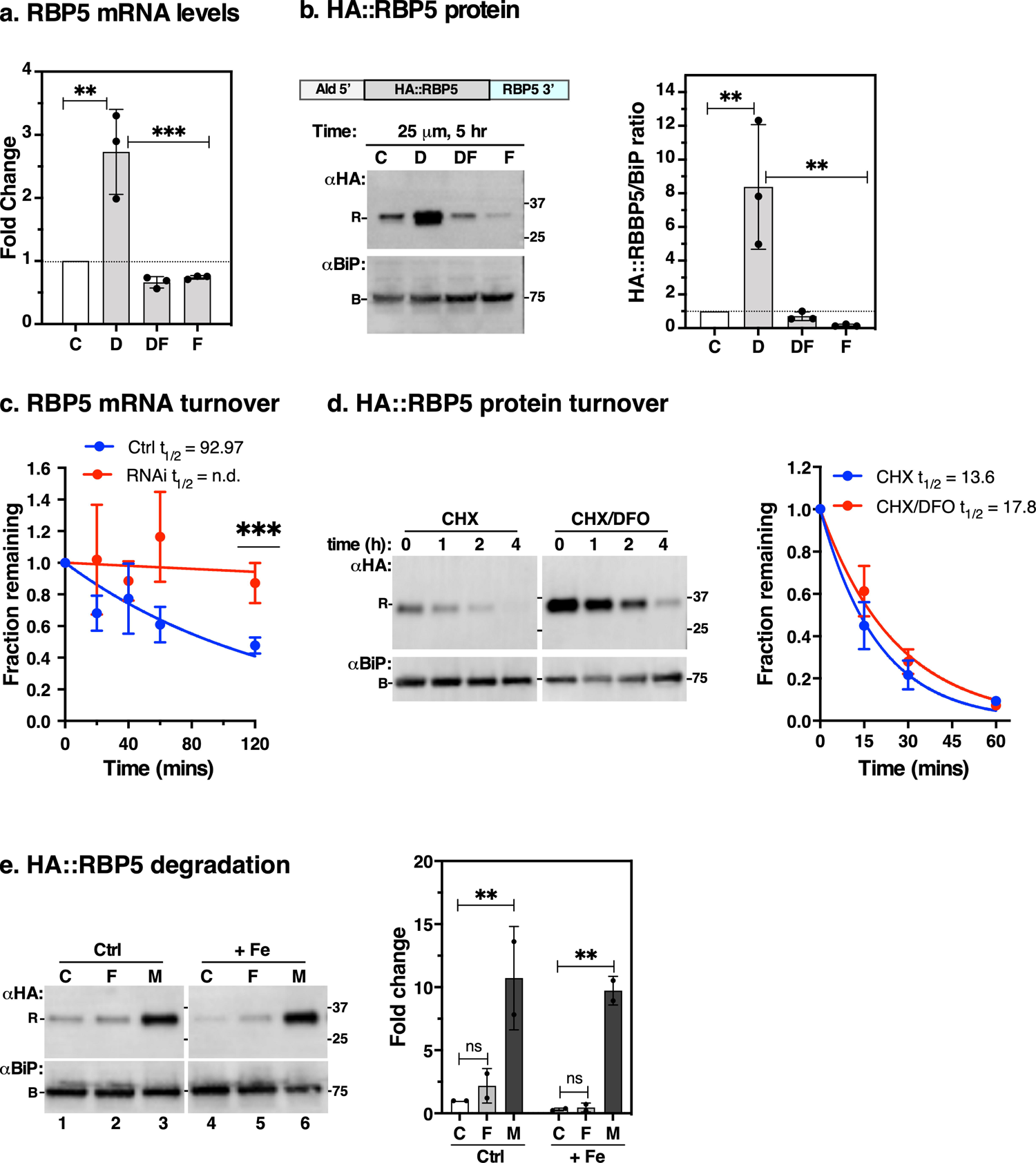
Effect of iron starvation on RBP5 stability. **a.** qRT-PCR analyses of endogenous *RBP5* mRNA levels following treatment of wild type cells as described in Figure 1. Data are means ± SD (*n* = 3 biological replicates; with 3 technical replicates for each *n*). *\*\*P* < 0.002, \**\*\*P* < 0.001, as determined by one-way ANOVA with post-hoc Tukey test. Only statistical comparisons with *p*-values < 0.05 are displayed. All others are not significant. **b.** Schematic of N-terminal *in situ* tagged RBP5 (HA::RBP5) preserving its native 3’-UTR – see *Supplementary Fig. S3*. HA::RBP5 cells were incubated with DFO as described above, total protein was immunoblotted with anti-HA (RBP5, R) or αBiP (B, loading control) antibodies. Bar chart (right) shows signals of HA::RBP5 from immunoblot quantified by densitometry using Image Lab software (Bio-Rad) normalised to the corresponding BiP signals relative to control samples. Data are means ± SD (*n* = 3 biological replicates). **c.** Endogenous *RBP5* mRNA turnover. The half-life of *RBP5* was determined by qRT-PCR using cDNA samples derived from TfR knockdown (RNAi) or control (Ctrl) cells incubated with Sinefungin and Actinomycin D as described above. Data are means ± SD (*n* = 4 biological replicates; with 2 technical replicates for each *n*). \**\*\*P* < 0.001 as determined by multiple unpaired t-tests. **d.** HA::RBP5 cells were incubated with 100 μg/ml cycloheximide (CHX) to block protein synthesis following pre-treatment with DFO (CHX/DFO, 5 hr) or without (CHX). Protein levels were analysed at time intervals indicated by immunoblotting with αTfR or αBiP (loading control). Decay curve shows rate of HA:RBP5 protein turnover quantified from immunoblots and normalised to the corresponding BiP signals at indicated time points relative to that of time zero. CHX and CHX/DFO blots were digitally separated for presentation post acquisition. Blot is representative of *n* = 3 biological replicates and plot shows means ± SD. **e.** RBP5 stability. HA::RBP5 cells were incubated with (+ Fe) or without (Ctrl) 25 μM FeCl_3_, 5 hr) supplemented with FMK024 (F, 20 μM) or MG132 (M, 25 μM) or vehicle (C) as indicated. Samples (∼5 x 10^6^ cell equivalents/lane) were collected and immunoblotted with anti-HA (αHA) or anti-BiP (αBiP, loading control). Bar chart (right) shows signals of HA::RBP5 quantified by densitometry using Image Lab software (BioRad). HA:RBP5 signal is normalised to the corresponding BiP signal and expressed as a ratio relative to lane 1. Fe (+) and Ctrl blots were digitally separated for presentation post acquisition. Blot is representative of *n* = 2 biological replicates and plot shows means ± SD. *\*\*P* < 0.002, as determined by two-way ANOVA with post-hoc Tukey test.

To test whether the *RBP5* 3’-UTR contributed to its regulation, we fused it with an mNG::Ty epitope tag *in situ* at the C terminus (37). Here, we replaced *RBP5* endogenous 3’-UTR with Paraflagellar Rod (PFR) 3’-UTR but retained its native 5’-UTR (**Fig S4b**). This modification abolished sensitivity of RBP5::mNG::Ty protein to iron when incubated with DFO (**Fig S4c**).

Northern blot analyses using a probe to RBP5 ORF and its UTRs showed a ∼4-fold increase in total mRNA abundance in DFO-treated samples relative to controls, with a single homogenous product of ∼1.9 kb in length (**Fig S4d**). Collectively, these results indicate that native *RBP5* 3’-UTR is necessary to confer an iron response and no additional RNA processing occurs in DFO-treated conditions relative to controls.

To determine the level of post-transcriptional regulation of *RBP5,* we first attempted to measure its mRNA half-life in the presence and absence of DFO. We found no significant difference in DFO-treated (t_1/2_ = 166.6 min) compared to controls (t_1/2_ = 152.3 min), although our half-life values were similar to previously reported values (35). However, qRT-PCR analyses showed that *RBP5* mRNA was significantly more stable in TfR-silenced cells compared to controls (**Fig 6c**), suggesting that changes in *RBP5* mRNA stability mediate iron responsive changes in its steady state levels. Since RBP5 protein stability remained unchanged compared to controls (**Fig 6d**), we conclude that increased mRNA stability contributes to iron responsive changes in RBP5 protein levels.

Our finding that RBP5 protein significantly decreased (*p < 0.05*) in iron replete conditions (**Fig 6b**, compare lanes F vs C) prompted us to investigate if this was due to increased cellular degradation. We incubated HA::RBP5 cells with or without FeCl_3_ in the presence of FMK024 or MG132 to block lysosomal or proteasomal degradation respectively. Without any protease inhibitors, the steady state level of HA::RBP5 decreased during 5 hr incubation with FeCl_3_ (**Fig 6e**, compare lanes 1 and 4), consistent with **Fig 5b**. FMK024 treatment was unable to rescue HA::RBP5 turnover (compare lanes 1 & 2; 4 & 5). Contrastingly, under normal or excess (+ Fe) iron, MG132 dramatically blocked HA::RBP5 turnover (compare lanes 1 & 3; 4 & 6), consistent with degradation in the proteasome. Collectively, these data indicate that when iron is sufficient RBP5 is highly unstable and rapidly degraded by the proteasome.

### RNAi and knockout suggest that *RBP5* is essential

To study the function of RBP5, we created a tetracycline inducible *RBP5* RNAi cell line using the plew100v5x stem-loop vector (38). RNAi for 72 hr showed no growth defect despite consistently observing a ∼90% reduction in *RBP5* mRNA (**Fig S5a & S5b**). To assess if RBP5 is required for *TfR* mRNA stability during iron starvation, we induced *RBP5* RNAi for 20 hrs followed by DFO treatment for 5 hr. We found ∼90% reduction in *RBP5* mRNA as judged by qRT-PCR, indicative of efficient depletion. However, the effect of DFO on *TfR* transcript stability was unaffected in the presence or absence *RBP5* depletion (**Fig S5c**), suggesting that partial loss of *RBP5* had no effect on iron-dependent regulation of *TfR*. In the absence of RBP5 antibody, the extent of RNAi on RBP5 protein depletion was assessed by transfecting the RNAi construct into HA::RBP5 expressing cells. Following 24 hr RNAi induction only, residual RBP5 protein was still detected by western blot (**Fig S5d),** indicating a direct correlation between loss of mRNA and protein.

To completely eliminate all RBP5 activity, we attempted to delete both alleles of *RBP5* by replacing the ORF with drug selection cassettes. First, using a conventional knockout strategy by cloning ∼400 nts of the intergenic regions flanking drug selection cassettes, whenever both drug selection markers were integrated, a third RBP5 allele had been duplicated in the genome, as judged by PCR (**Fig S5e)**. In two independent transfections we did not obtain any viable clones. Second, conditional knockouts attempts where an ectopic copy driven by inducible doxycycline allows replacement of the second allele were also unsuccessful because RBP5 overexpression is toxic (see below). Collectively, these observations argue that BSF *T. brucei* cannot tolerate complete loss of *RBP5* and that extremely low levels of RBP5 protein are required to maintain cell viability.

### Elevated RBP5 levels disrupt normal cell cycle progression

To determine if RBP5 overexpression affects cell fitness we generated cells harbouring an ectopic tetracycline inducible *RBP5* copy, leaving both endogenous *RBP5* alleles intact. Treatment with 1 μg/ml tetracycline resulted in ∼15-fold increase in *RBP5* mRNA as judged by qRT-PCR (**Fig 7a**) and this reduced cell growth after 3 days of overexpression (**Fig 7b**). To distinguish between acute lethality and reduced growth rates, we performed a live/dead assay by incubating live cells with propidium iodide (PI). Following 24 hr of overexpression, while uninduced cells remained PI-negative (alive), the mean fluorescence intensity of induced cells increased ∼5-fold as indicated by a shift to the right in fluorescence (**Fig 7c, left**). We interpret these to be cells where the membrane integrity was compromised, and thus permeable to PI. **Fig 7c, right and 7d** show the DNA content of fixed PI- and DAPI-stained trypanosomes analysed by flow cytometry and microscopy, respectively. The flow cytometry data showed that RBP5 overexpression resulted in an increase in the sub-G1, S and G2/M peaks (**Fig 7c, right**). Analyses of DAPI-stained cells (**Fig 7d**) showed a reduction in cells with one nucleus and one kinetoplast (1K1N) by 20% and simultaneous increase in cells with 0N1K and aberrant N-K configurations by 20%, in agreement with flow cytometry data. Taken together, we conclude that excess RBP5 of ∼10-fold greater than observed during iron starvation selectively disrupts normal cell cycle progression and hence normal cell growth.

**Figure 7.**
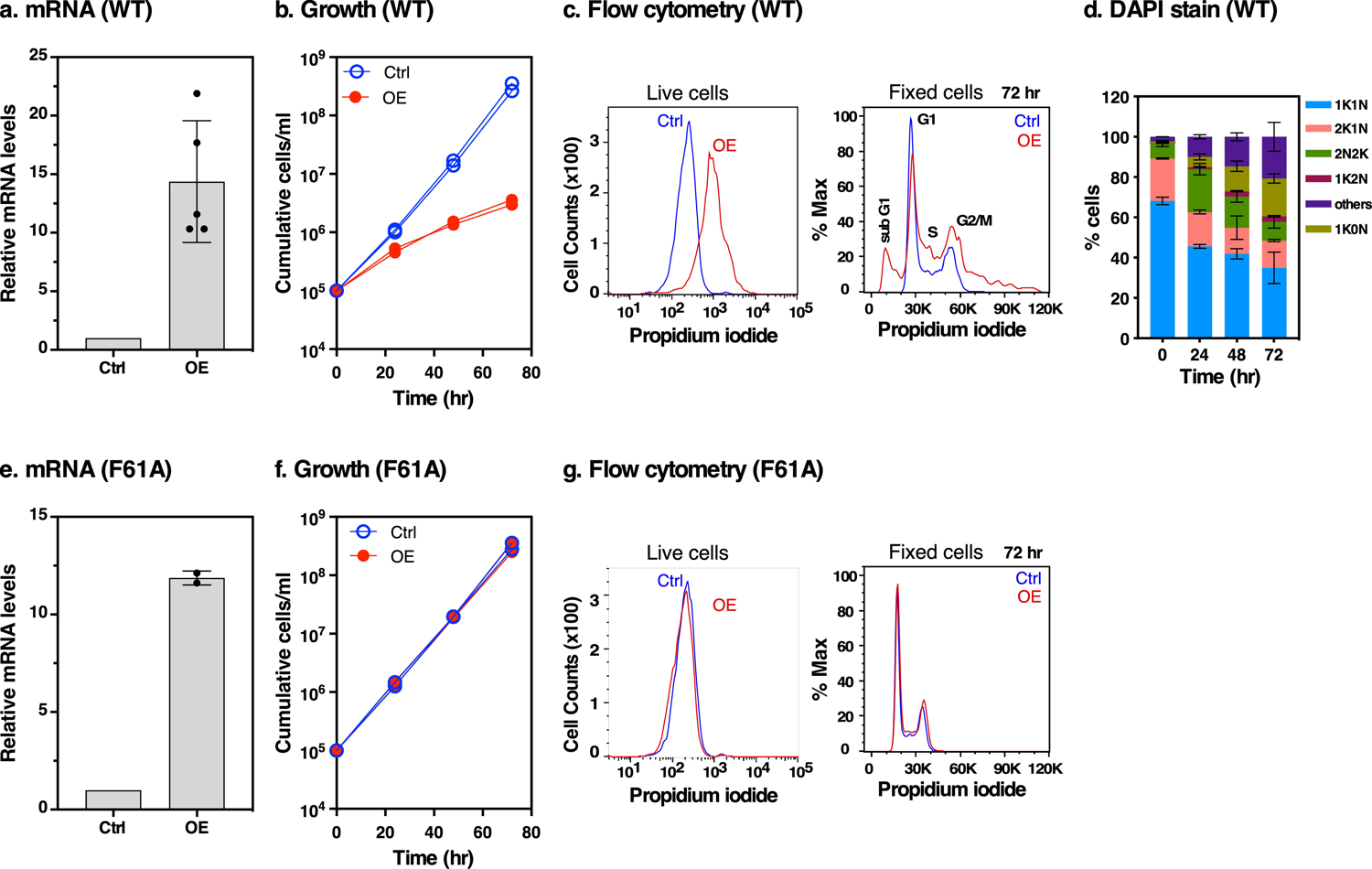
Effects of *RBP5* wild type (WT) and F61A mutant overexpression. **a, e**. qRT-PCR analyses of *RBP5* mRNA levels following 24 hr overexpression of wild type *RBP5* (WT) or *F61A* mutant with (OE) or without (Ctrl) 1 µg/ml tetracycline. Phe at position 61 is mutated to Ala in F61A. Data are means ± SD (*n* = 5 biological replicates for WT and n = 2 for F61A; with 3 technical replicates for each *n*). **b, f.** Effect of *RBP5* overexpression on cell growth. Induction of RBP5 WT or F61A mutant was achieved with (OE) or without (Ctrl) 1 µg/ml tetracycline. The y-axis shows cumulative number of parasites/ml enumerated using a hemocytometer. **c, g.** Analysis of cell membrane integrity and DNA content by flow cytometry. After 24 hr of incubation with tetracycline (OE) or without (Ctrl), live cells (left) were incubated with propidium iodide as described in methods. In both cases single trypanosomes were gated by forward and side scatter to exclude doublets and cell debris. 10,000 trypanosomes were analysed per sample as shown on the y-axis as counts (live). The total number of gated single cells were equivalent in both OE and Ctrl live cells. For fixed cells, the total number of gated singlets were less than 20% of Ctrl cells hence normalisation to % Max. Note the appearance of cells with sub-G1 DNA (peak at far left) following OE, as well as cells with higher order DNA content (far right). Flow cytometry histograms are representative of two biological replicates of two independent clones. Cell cycle phases G1 and G2/M are indicated representing cells with chromosomal content 2C and 4C, respectively. **d.** Following RBP5 overexpression at the time points indicated on the x-axis the number of nuclei (N) and kinetoplasts (K) per cell (n > 200) were microscopically analysed. Cells with 1N1K are in G1 or S phase, cells with 1N2K are in G2, and cells with 2N2K are mitotic or post-mitotic. Cells with aberrant number of kinetoplasts or nuclei (0N0K or >2N2K) were scored as others.

To test if the lethality of RBP5 overexpression was due to its RNA binding activity, we mutated Phe at position 61 to Ala (F61A), maintaining hydrophobicity at this position but potentially disrupting the most conserved RRM signature motif RNP1 [RK]-G-[FY]-[GA]-[FY]-[ILV]-X-[FY] in the β3 strand (39) (**Fig S6)**. Three days of F61A overexpression resulted in ∼12-fold increase in mRNA levels (**Fig 7e**) but had no effect on growth (**Fig 7f**) or cell membrane integrity (**Fig 7g**), providing evidence that the growth defect of wild type overexpression was due to its RNA binding activity.

### RBP5 associates with its own and PAP2 mRNAs

In order to identify mRNAs that associate with RBP5, HA::RBP5 expressing cells were first treated with or without DFO for 5 hr. Next, pull downs were performed with anti-HA antibodies followed by RNA isolation and qRT-PCR (RIP-qRT PCR) analyses (**Fig 8a**). Anti-HA immunoblotting showed more pulldown of RBP5 in DFO-treated cell lysates relative to controls (**left**) and specific enrichment of *RBP5* (∼6-fold), *PAP2* (∼2-fold) and Hyp#2 (∼2-fold) mRNAs (**right**). We found no enrichment of TfR transcripts (*E6, E7*), other iron regulated mRNAs identified by RNA-Seq, or unrelated (28s rRNA and GAPDH) mRNAs which served as non-specific controls. To test if RBP5 association with these mRNAs was dependent on its RNA binding activity, we performed RIP-qRT PCR on tetracycline inducible Ty-tagged wild type RBP5 using the non-RNA binding F61A mutant as control (**Fig 8b**).

**Figure 8.**
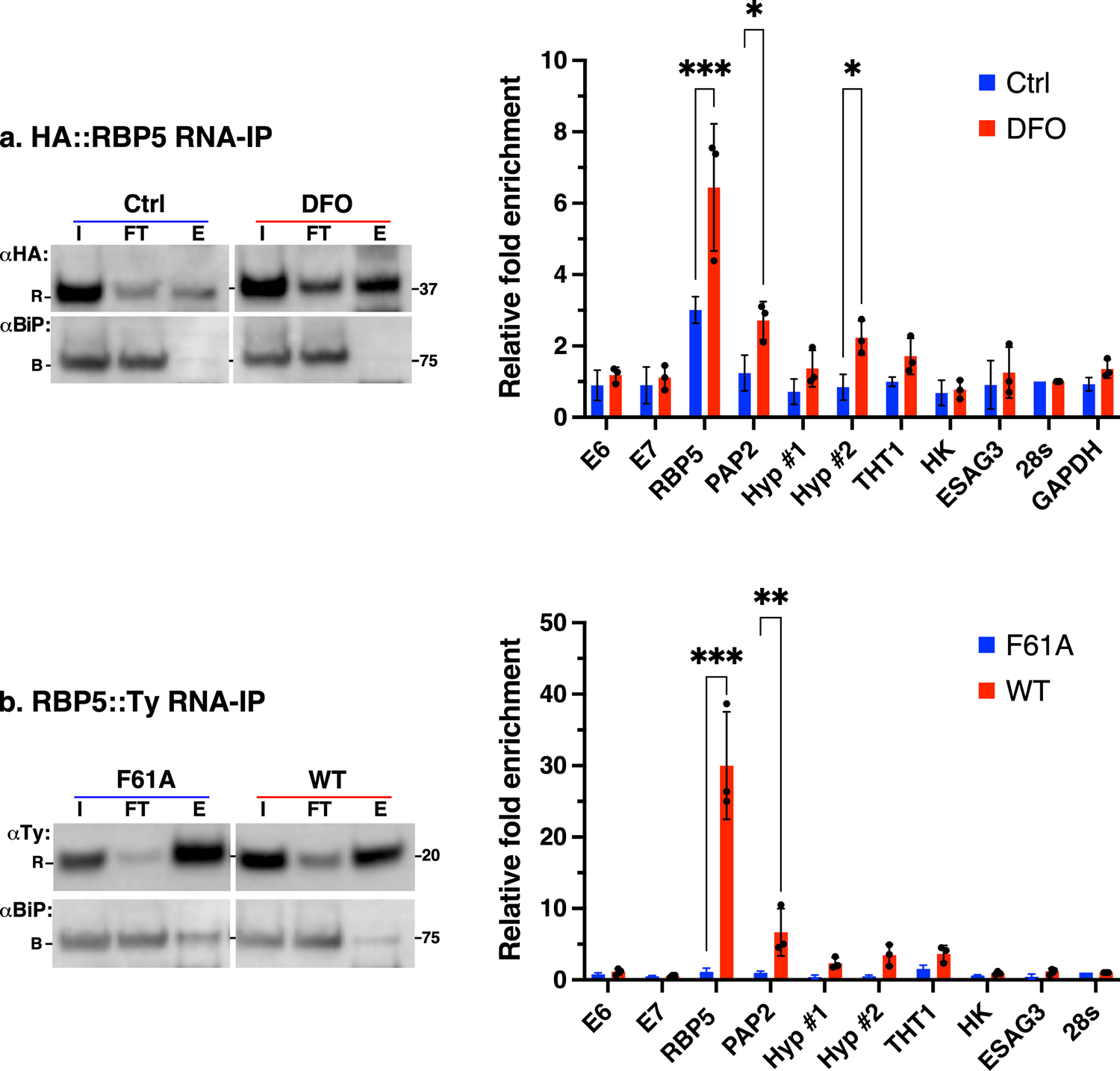
Identification of RBP5-associated mRNAs. **a.** HA::RBP5 cells were incubated without (Ctrl) or with (DFO) deferoxamine (5 hr) and RBP5 pulldowns were performed with anti-HA beads. **Left**: Western blot analyses of input (I), flow through (FT) or Eluate (E) fractions with antibodies for HA::RBP5 (αHA, R), and BiP (αBiP, B). **Right**: RNA was purified from “FT” (input) and “E” (IP) fractions and qRT-PCR performed as described above. Bar chart shows relative fold-enrichment of mRNAs shown on the x-axis as determined by the equation 2 *^Ct^*, where *C_t_* = *C_t_* (input) − *C_t_* (IP), C*_t_* = cycle threshold, normalised to 28s rRNA as non-bound control. Data are means ± SD (*n* = 3 biological replicates: with 2 technical replicates for each *n*). \**\*\*P* < 0.001, *\*\*P* < 0.002, *\*P* < 0.05 as determined by two-way ANOVA with post-hoc Sidak test. Only statistical comparisons with *p*-values < 0.05 are displayed. All others are not significant. **b.** C-terminally Ty-tagged wild type RBP5 (WT) or an RRM mutant (F61A) was immunoprecipitated with anti-Ty beads from overexpressing BSF cells simultaneously treated with tetracycline and DFO for 4 hours. **Left:** RNPs were isolated as described in (a) above and blotted with anti-Ty antibodies for WT or F61A (αTy, R) and BiP (αBiP, B). **Right:** Bar chart shows relative fold-enrichment of transcripts shown on the x-axis. Fold enrichment and normalisation were performed as in (a). Note: To assess if DFO treatment had an effect on RBP5 mRNA enrichment, the experimental setup was repeated without or with DFO and the data are shown in supplementary **Fig S7**.

Blotting with anti-Ty antibodies confirmed efficient pulldowns of both WT and F61A mutant (**left**), with specific enrichment of only *RBP5* (∼25-fold) and *PAP2* (∼5-fold) transcripts confirming their interactions with WT but not with F61A (**right**). We found no enrichment of *RBP5* mRNA when RIP was performed with untagged parental or F61A cell lysates establishing the specificity of these interactions (**Fig S7**). Taken together, these experiments only allow us to conclude that increased RBP5 expression either by DFO treatment (5 hr) or moderate overexpression (4 hr) results in specific interactions with its own and *PAP2* transcripts. The observed difference in mRNA fold enrichment between DFO treatment and overexpression reflects the amount of available protein. It is unclear why Hyp#2 mRNA was enriched in DFO-treated but not with overexpression.

### RBP5 and TfR upregulation occur simultaneously during iron starvation, preceding PAP2 activation

To determine the kinetics of the iron response, we performed a time course experiment in which cells were starved of iron either by DFO or TfR silencing. qRT-PCR showed that native *E6* and *RBP5* mRNA were elevated within 60 min of DFO incubation and increased exponentially as a function time (**Fig 9a**). The same pattern was observed in TfR knockdown cells (**Fig 9b**), albeit with a 5-hr lag due to the time required for TfR ablation. The disappearance of native *E7* mRNA coincided with accumulation of *RBP5* and RNAi-resistant *E6^R^* transcripts starting at 8 hr post RNAi induction and plateauing at 24 hr.

**Figure 9.**
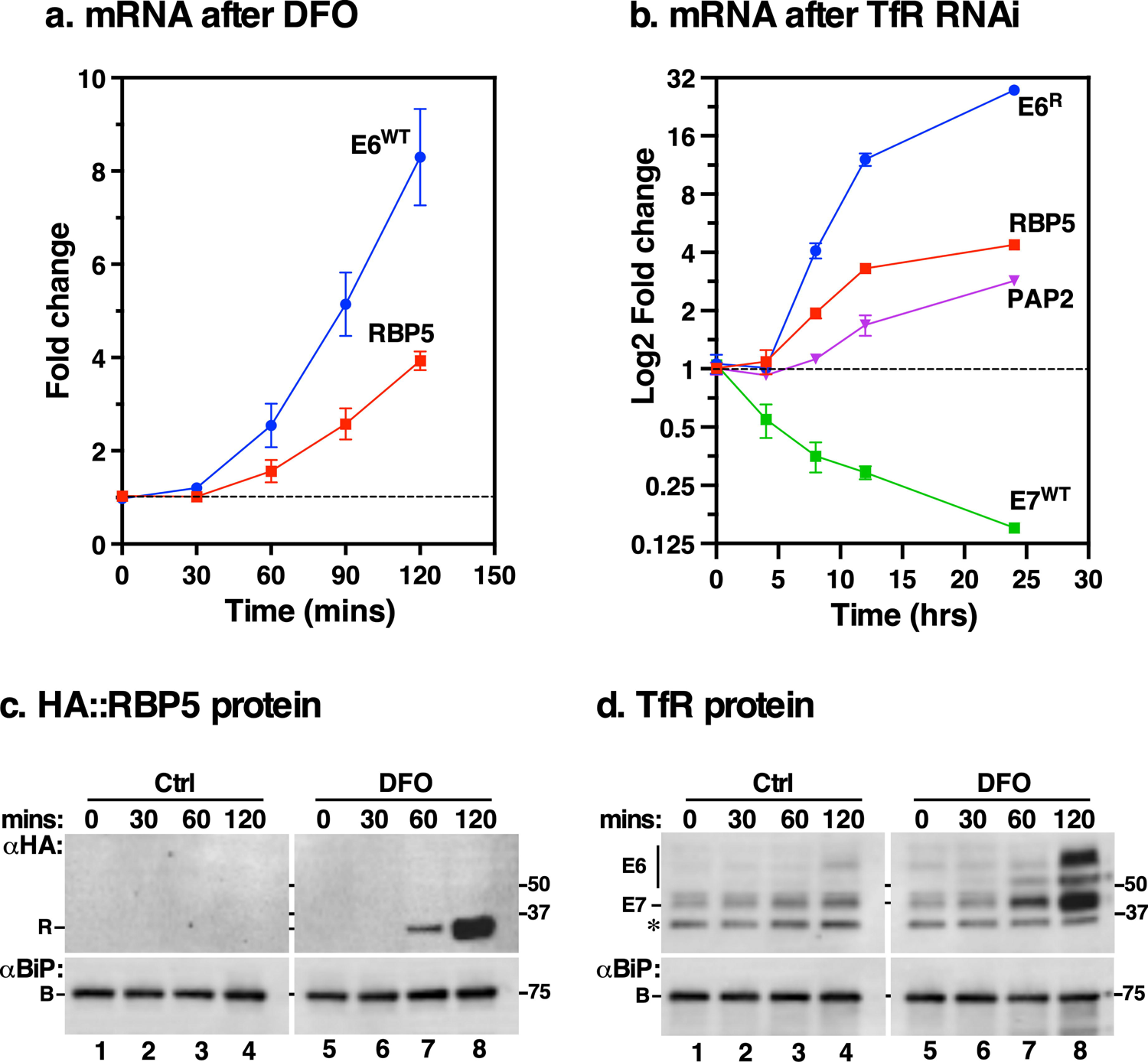
Time-course response of TfR, RBP5 and PAP2 expression to iron starvation. **a.** Determination of the iron starvation response kinetics. Wild type cells were treated with DFO for 120 mins, total RNA harvested at time points shown on the x-axis, and relative mRNA levels determined by qRT-PCR for indicated genes (inset). The rate of accumulation of native *E6* (*E6^WT^*) and *RBP5* signals is normalised to the equivalent signal from un-induced time zero cells relative to *ZFP3* as endogenous control. Data are means ± SD (*n* = 3 biological replicates; with 3 technical replicates for each *n*) **b.** Following native TfR silencing for 24 hr in E6^R^ expressing cells, total RNA was isolated at specified time intervals (x-axis) and mRNA levels of TfR (*E7^WT^*, *E6^R^*) *PAP2,* and *RBP5* were analysed by qRT-PCR. Normalisation followed the same procedure in (a). Data are means ± SD from (*n* = 3 biological replicates). **c.** Kinetics of HA::RBP5 protein. HA::RBP5 cells were treated with DFO for 120 min, total protein was extracted at indicated time points and blotted with either anti-HA (R) or anti-BiP (B, loading control) antibodies. Note: image acquisition was performed maintaining a linear dynamic range across all time points. The low HA::RBP5 signal in Ctrl blots is masked by the intense signal at 120 min time point. Ctrl and DFO blots were digitally separated for presentation post acquisition. Blot is representative of three independent biological replicates. **d.** Kinetics of TfR protein. Samples from (c) above were blotted with either anti-TfR (E6, E7) or anti-BiP (B, loading control) antibodies. Blot represents *n* = 3 as in (c). Star indicates a non-specific background polypeptide.

Accumulation of *PAP2* transcripts, however, began at 12 hr, four hours after induction of *RBP5* and *TfR*. These data suggest that when internal iron pools are depleted, systemic iron responsive signals are activated within the first hour and the effects of such signals on *RBP5* and *TfR* transcripts occur simultaneously, preceding the PAP2 iron response. Finally, we monitored the time interval of RBP5 and TfR protein response and found that the earliest response with DFO occurred within the same time frame as their mRNAs (**Fig 9c and 9d**), suggesting that there is no time lag between *RBP5* and *TfR* mRNA and protein regulation.

## Discussion

Adaptive iron responses are critical for invading pathogens to maintain successful parasitism. While similarities exist in iron regulatory pathways between pathogens and host cells, substantial differences also prevail. In our search for iron regulatory factors in *T. brucei* by transcriptomics, we used two modes of iron starvation (DFO and TfR RNAi) known to elevate expression of endogenous TfR (30, 31). We find that beside TfR, BSF *T. brucei* upregulates a cohort of trypanosome-specific genes (ESAG3, PAGS, RBP5), genes involved in glucose uptake and glycolysis (THT1 and hexokinase), endocytosis (PAP2), and most notably an RNA binding protein RBP5 (**Fig 2)**. This finding demonstrates that post transcriptional regulation of iron responsive genes deviates from “standard” eukaryotic model systems (36), including closely related *Leishmania* and host cells. The significance of each of these factors is discussed below.

First, we show that iron depletion by DFO inhibits growth of BSF *T. brucei in vitro* while free iron [25 μM] shows negligible toxicity (**Fig 1**), consistent with previous work (32). Although physiological concentrations of free iron in the host never exceed 1 μM (40) due to associated toxicity, certain pathologies can result in free iron concentrations as high as 20 μM (40), similar to levels used in our experiments. Moreover, Breidbach *et al* incubated BSF parasites at concentrations 4-fold greater (100 μM ferric ammonium citrate) and showed that it rescued the inhibitory effects of DFO (32), suggesting that BSFs tolerate intermittent high levels of free iron. Such levels can be experienced during the chronic anaemic phase of trypanosome infections reported both in natural and experimental scenarios, characterised by increased iron release for erythropoiesis and high parasitaemia (41), or in the insect midgut right after a bloodmeal where significant breakdown of heme occurs. In closely related *Leishmania amazonensis*, iron toxicity is alleviated by LIR1, a plasma membrane iron exporter that prevents excessive intracellular accumulation of labile iron (42). A LIR1 orthologue is present in *T. brucei* and may fulfil the same role, but this remains to be shown. Second, prolonged iron starvation by TfR silencing corresponds to situations where limited iron availability might be encountered such as during skin and adipose tissues colonisation (1, 2) where Tf concentrations are negligible, in *T. vivax* where no TfR related genes appear to be present (43), or during transitions from the blood to insect vectors where TfR expression is transcriptionally silenced.

The mechanisms of iron-dependent regulation of TfR expression has remained elusive for 20 years (31). Upregulation of TfR is mediated, at least in part, through unidentified *cis* regulatory elements encoded in its 3’-UTR but aconitase is not the *trans-*acting factor (31, 33). We show that the *TfR* 3’-UTR is sufficient to mediate stability of a reporter construct, consistent with a previous study (33). Our finding that protein stability is not a contributing factor adds a new dimension (**Fig 3**). In the absence of post-translational control, *T. brucei* adaptation to iron deprivation is driven by increased *TfR* mRNA stability due to reduced turnover. Stability is likely conferred either by mRNA recruitment to the translation machinery as previously suggested or through an RBP that binds to the 3’-UTR (44), or both. Increased *TfR* mRNA stability therefore accounts for elevated protein levels, ultimately working to boost efficient iron import to maintain homeostasis.

In contrast to upregulating TfR expression levels to maintain iron homeostasis, we find that free iron reverses the TfR RNAi growth defect and abolishes intracellular iron response signals. These observations argue that alternate iron uptake occurs in *T. brucei.* The simplest explanation is acquisition of free iron by fluid phase endocytosis, followed by reduction and transport in the endo/lysosomal compartments by Mucolipin (13). Reports of extremely high endocytosis rates in BSF *T. brucei* (45) and of both receptor-mediated and fluid-phase endocytosis occurring through the flagellar pocket (46, 47) argue that fluid-phase uptake provides a passive iron entry route when extracellular free iron is elevated. However, growth defects observed in TfR silenced cells suggest that fluid-phase endocytosis of holo-Tf alone is insufficient to provide enough iron for optimal growth under normal culture conditions. Alternatively, exogenous iron may enter BSF cells through dedicated surface localised machinery that reduce Fe^3+^ and import Fe^2+^ directly into the cytosol. Such a mechanism was reported for PCFs and is backed by the presence of two putative iron reductases and iron transporters in the *T. brucei* genome whose function and location remain unknown (20). This two-step reductive mechanism has been described for *S. cerevisiae* (48), *Naegleria fowleri* (49) and the closely related Kinetoplastid *Leishmania. Leishmania* ferric reductase 1 (LFR1) reduces Fe^3+^ to Fe^2+^ which is subsequently transported to the cytosol by *Leishmania* iron transporter 1 (LIT1) (50, 51). Orthologues of both are present in the *T. brucei* genome (25). We predict that endocytosis of holo-Tf and reductive iron import work together to satisfy parasite iron needs during infection. Such a mechanism could be operational in *T. vivax* where comparative genomics data show that this species lacks VSG-like TfRs (43).

An interesting finding was upregulation of phosphatidic acid phosphatase superfamily enzymes: PAP2 (Tb927.8.480, Tb08.27P2.110), which was abolished by functional complementation (**Fig. 4**). In eukaryotes, PAP enzymes regulate lipid synthesis, catalysing Mg^2+^-dependent dephosphorylation of phosphatidic acid to diacylglycerol (DAG) (52), a second messenger for intracellular signalling (53). In *T. brucei*, genetic and small molecule experiments showed that DAG produced by GPI-PLC stimulates Tf/Fe endocytosis (53). PAP2 in *T. brucei* is a transmembrane protein and our preliminary experiments showed distinct puncta between the nucleus and kinetoplast indicative of endosomal localisation (Cornel, L and Tiengwe, C; unpublished). Under conditions of low TfR expression, upregulation of PAP2 may therefore boost DAG production stimulating Tf/Fe endocytosis by fluid phase.

RBP5 is the most significant of the proteins upregulated by iron starvation. Like TfR, its expression is modulated by iron levels (**Fig 6**) in both DFO-treated and TfR silenced cells. RBP5 protein has a single RRM domain in its N-terminus, is cytosolic, and is conserved in trypanosomatids, indicative of an evolutionarily common function. RBP5 stability is very atypical: the mRNA is extremely stable (t_1/2_ > 1.5 h) while the protein is remarkably unstable (t_1/2_ = 13.6 min). The median half-life of mRNAs and proteins in BSF *T. brucei* is 12 min and 5.6 hr respectively (35, 54). Normal turnover of RBP5 is mediated by the proteasome since the protein abundance increases with prior incubation with MG132, suggesting that it is unstable. RBP5 post-translational stability is unaffected by iron levels. Thus, its 3’-UTR primarily controls iron responsive changes in mRNA levels.

To our knowledge, RBP5 is the first iron responsive RNA binding protein identified in BSF *T. brucei*. Available transcriptome data show that it is significantly upregulated when trypanosomes experience iron scarce environments. For example, in adipose tissues relative to the bloodstream (2) or in the salivary glands relative to the midgut during development in the tsetse fly vector (55). Bioinformatic analyses suggest that the protein lacks putative iron binding motifs. Therefore, increased *RBP5* mRNA abundance (and consequently protein) is mediated by a feedback mechanism via direct nucleotide interactions with a *cis* IRE in its 3’-UTR or via accessory trans-acting factor(s). The actual *cis* element, associated factor(s) and mechanisms remain to be determined. Unlike TfR, however, we cannot rule out a translational effect because we observe a maximum 4-fold increase in RBP5 mRNA abundance but an 8-fold increase in protein levels. mRNA stability therefore accounts for most, but not all increase in RBP5 protein levels. During iron deficiency, association of RBP5 with its mRNA results in increased cytosolic protein concentration which in turn interacts with *PAP2* mRNAs downstream. The simplest explanation for PAP2 upregulation is increased DAG production stimulating endocytosis (53). Data showing that PAP2 is activated 4 hrs after RBP5 upregulation (**Fig 9**) and that there is significant increase in endocytosis of tomato lectin in TfR-deficient cells (**Fig S1**) support this model. Nevertheless, our results show that increased RBP5 expression in iron-deprived cells has a maximum threshold because overexpression above this threshold (∼15 fold) is toxic.

Toxicity associated with RBP5 over-expression can be attributed to its RNA binding activity conferred by the RRM (**Fig. 7)**. Alignment of RBP5 RRM with RRM1-family proteins from PFAM suggests a close relationship with proteins from *Drosophila*; *Homo sapiens*; *S. cerevisiae* and *Schizosaccharomyces pombe* (**Fig S6**). These proteins have multiple RRMs that arose from duplication events providing specificity for mRNA targets, additional functionality of intra- and/or inter-protein:RNA or protein:protein interactions (56). A single RRM is usually insufficient to define specificity of RBPs (57), yet RBP5 has just one RRM, suggesting that homo- or heterotypic oligomerisation(s) are important for defining target specificity and function. In *Drosophila* ELAV and SXL proteins, aromatic mutations in the RNP1 domain resulted in a non-functional RRM (56). Similarly, the null phenotype of our F61A overexpression cells (one of the most conserved residue in RRMs) supports a potential disruption of RNA-RBP5 interactions impairing its RNA binding activity. RBP5 toxicity is associated with disruption of normal cell cycle progression caused by an accumulation of anucleate cells and cells in G2/M phase. Whether this implicates RBP5 (directly or indirectly) in regulation of cell cycle related genes remains to be determined.

Finding other RBP5 mRNA targets will be crucial for understanding if it has additional roles in cell cycle. In the absence of its global associated factors, the evidence presented form an interesting basis for discovery of novel post-transcriptional mechanisms coordinating iron homeostasis and cell cycle control. This connection is well documented in other systems (58, 59), but never reported in trypanosomes.

Knocking out both alleles of RBP5 consistently resulted in gene duplication producing a third ectopic allele. RNA silencing did severely reduce mRNA and protein (∼90%), but had no effect on growth, suggesting that extremely low levels are sufficient for survival. Knockdown in the presence of DFO did not reverse iron dependent *TfR* mRNA stability. These observations argue that either residual RBP5 protein is sufficient to maintain TfR stability, there is possible functional redundancy with an unknown gene, or RBP5 does not mediate TfR stability. Collectively our data favour the latter possibility because pulldown assays show no association of *TfR* mRNAs with RBP5 protein. Second, the mode of induction of both RBP5 and TfR in iron depletion is similar, increased mRNA stability and protein abundance. Third, the kinetics of RBP5 and TfR iron response is similar suggesting coordinate regulation, likely mediated by a common factor albeit yet undiscovered. It is possible that regulation of the unidentified TfR stabilising factor occurs by post-translational mechanisms precluding its identification by transcriptomics. Its regulation may also be mediated by non-coding RNAs (60) or by iron sensing riboswitches that directly bind iron (61). Experiments to test these possibilities will reveal insights on how *TfR* mRNA is stabilised in iron starvation.

Other iron responsive genes identified were ESAG3 and two hypothetical proteins (Tb927.8.490 and Tb927.8.510). Tb927.8.490 and Tb927.8.510 have predicted multi-pass transmembrane domains and/or signal sequences, predicted ion binding motifs, located within the secretory pathway compartments (http://tryptag.org/), suggesting they may either be secretory signalling effectors or execute ion transport functions. While ESAG3 protein location and function are unknown, it is essential for survival of BSF *T. brucei* (62). Whether it fulfils a role in iron homeostasis remains to be determined but its genomic location at BES suggests a host adaptation function like other ESAGs.

Paradoxically, iron starvation response resulted in upregulation of Procyclin-associated genes (PAGS), genes not expressed in BSF trypanosomes. Phylogenetic analyses of PAG and PAG-like genes in *T. brucei* and *T. congolense* suggest that they encode transferrin receptor-like proteins, often co-located in pairs with predicted GPI-minus and GPI-positive signals reminiscent of TfR subunits (43). The simplest interpretation of PAG upregulation is that some PAG genes may encode surface localised, yet uncharacterised, receptors that serve procyclic cells where TfR is transcriptionally silenced (63). Further experiments will be needed to prove this since PCFs do not bind transferrin.

In conclusion, our work has uncovered an unusual means by which BSF *T. brucei* reorganise their transcriptome to deal with iron stress. The pathway contrasts that of closely related *Leishmania* parasites which regulate transcript levels of iron transport proteins (*LFR1* and *LIT1*), like host cells (51, 64). We have defined the kinetics and mode of regulation of at least two early factors (TfR and RBP5) activated during iron starvation in BSF *T. brucei* and show that RBP5 protein interacts with its own and *PAP2* mRNAs. **Figure 10** summarises iron responsive processes following iron starvation by TfR depletion. Although TfR is essential for optimal growth *in vitro* (29, 30), our data argue that it is not absolutely necessary for survival, suggesting that any therapeutic considerations for TfR must be evaluated with caution, bearing in mind that alternative iron entry systems prevail. In contrast, RBP5 expression level is important for cell viability. Its co-regulation with Hexokinase and THT1 hexose transporter in TfR-depleted cells suggests it might also have broader roles in general nutrient sensing. For example, it has been reported to be upregulated in glucose-starved cells (65), a finding that we have reproduced experimentally (C. Tiengwe, unpublished). Identifying its mRNA targets globally and molecular partners will be important for revealing its cellular dynamics.

**Figure 10.**
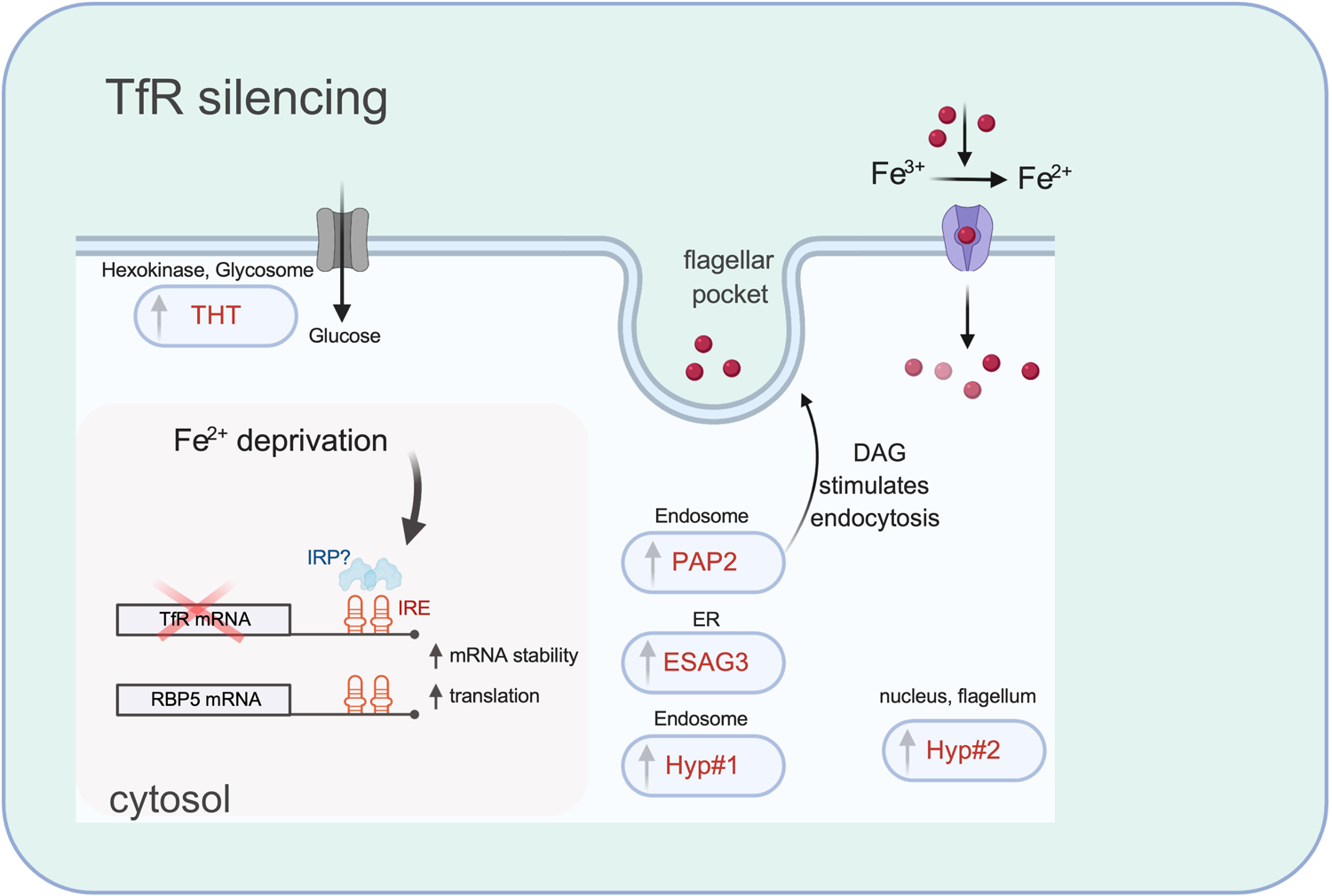
Illustration of the main iron responsive regulatory machinery of BSF *T. brucei*. Transcriptome and qRT-PCR data (**Figs 2, 3, 6, & 9**) from TfR depleted and DFO-treated cells show increased steady state levels of *RBP5* and *TfR* mRNA and protein. **Figs 2, 6 & S4** show that the 3’-UTRs of *TfR* and *RBP5* mRNAs confer stability suggesting they contain an iron responsive element (IRE) acting in *cis*. Figs 2 and 6 show that protein stability does not contribute to TfR and RBP5 iron starvation response indicating that increased translation drives elevated protein levels. Red cross depicts TfR depletion by RNAi. Upwards pointing arrows depict transcripts increased in TfR silenced cells. With exception of ESAG3 and PAP2, location of iron responsive proteins were derived from http://tryptag.org/. PAP2 and ESAG3 localisation are derived from our unpublished work (Cornel, L; Magnus, L; and Tiengwe, C; unpublished). It remains to be shown whether in the absence of TfR, free iron (red spheres) uptake occurs by fluid phase endocytosis or through a two-step reductive process. Image is created with BioRender.com.

## Materials and methods

### *T. brucei* BSF cell culture and iron starvation inductions

We used a tetracycline (Tet) inducible single-marker bloodstream form Lister 427 strain *T. brucei brucei* for all experiments (66). The cell line allows for Tet-regulated expression of transfected constructs for RNAi or overexpression achieved by addition of 1 μ cultured at 37°C, 5% CO_2_ in HMI-9 medium supplemented with 20% heat inactivated Foetal μg/ml streptomycin (Gibco). Cell lines were µg/ml G418 (parental) and 2.5 μ Phleomycin for inducible RNAi or overexpression cells, and Hygromycin (5.0 µg/ml), and Puromycin (0.25 µg/ml) for reporter or epitope-tagged cell lines. All iron starvation experiments were performed at cell densities ≤ 5×10^5^ cells/ml. Iron starvation was induced for 5 hr as follows: with 25 μM Deferoxamine (D, Sigma Aldrich), 25 μM Deferoxamine pre-saturated with 27.5 μM FeCl_3_ (DF), 25 μM FeCl_3_ (F, Sigma Aldrich); unless otherwise stated. Pan-specific RNA knockdown of TfR was induced for 24 hours. The constructs for TfR silencing, targeting and RNAi-resistant functional complementation have been previously published (30) and were a generous gift from Prof. J. D. Bangs (University at Buffalo, NY, USA).

### Quantitative real time PCR (qRT-PCR)

All qRT-PCR analyses used ∼1×10^7^ cells total and RNA was extracted from cell pellets using Qiagen RNeasy Extraction Kit. 500 ng RNA was used in cDNA synthesis using Biorad iScript cDNA synthesis kit per manufacturer’s instructions. cDNA was diluted 1:10 and analysed using Biorad iTaq SYBR Green on Applied Biosystems StepOne Real-Time PCR System (Life Technologies, Carlsbad, CA).

All reactions were performed in technical triplicates and specificity of each reaction was checked by ensuring single peaks from melt curves. Fold change was calculated by comparative C_T_ method using the Applied Biosciences software. Relative transcript levels were compared to endogenous control, *ZFP* (Tb927.3.720, nts 241-301) relative to untreated controls. Primer sequences are available in supplementary material.

### Library preparation and RNA-Seq analyses

Total RNA was harvested from 5×10^7^ cells total per condition, mRNAs were isolated from by oligo(dT) enrichment, first and second strand cDNA synthesized. cDNA fragments were purified for end reparation and single nucleotide A (adenine) addition. The cDNA fragments were linked with adapters and selected for PCR amplification. Agilent 2100 Bioanaylzer and ABI StepOnePlus Real-Time PCR System were used for quantification and qualification of libraries. Twelve samples (each sample: D, DF, F, and C were performed in technical triplicates) and sequenced on Illumina HiSeq platform, averagely generating about 10.39 Gb bases per sample.

Sequencing was performed by BGI technologies (Beijing, China). For TfR RNAi experiments, library Preparation and QC were performed by TruSeq RNA (Illumina). HiSeq 2500 Sequencing (50-Cycle single read sequencing per Lane. Total 6 samples: n = 3 technical replicates for TfR knockdown and 3 for controls. Sequencing was performed by UB Genomics and Bioinformatics Core (New York, USA).

### Bioinformatic analysis

Paired-end reads of biological replicates were aligned to the reference genome v46 of *T. brucei* clone TREU927 downloaded from TriTrypDB (67) using Bowtie2, with the ‘very-sensitive-local’ pre-set alignment option. The alignments were converted to BAM format, reference sorted and indexed with SAMtools. The genome coverage of the aligned reads was extracted from the BAM files using bedtools with the -bg option to output bedGraph files. The bedGraph files were visualized with the svist4get python package. Fragment counts were determined from the BAM files using featureCounts with parameters: -M (count multi-mapping) -O (match overlapping features) -t transcript (count level) -g gene_id (summarization level); for the paired end reads we added the parameters: -p (pair end) -B (both ends successfully aligned) -C (skip fragments that have their two ends aligned to different chromosome). Counts were performed using an in house developed GTF file that extends the annotation of the *T. brucei* 927 genes by including the VSG sequences deposited at http://tryps.rockefeller.edu/ and the Ksplice lncRNAs sequences (KS17_ prefix) described in (60). Finally we included newly predicted transcripts derived from a recent study aimed at characterising the full repertoire of expressed genes in *T. brucei* (Tinti *et al*., unpublished), using the Trinity (entries with TRY_ prefix) and Scallop (entries with MSTRG_ prefix) software.

### RNA-Seq quality control

The quality of alignments was evaluated with Qualimap2 using the bamqc and rnaseq options. The Qualimap2 output files, and the outputs of fastp, bowtie2, Picard Mark Duplicates, SAMtools flagastat, SAMtools stats and featureCounts were aggregated with MultiQC, and available at https://calvin-tfr.onrender.com/tfr_mqc/report.html for the DFO-treated samples and at https://calvin-tfr.onrender.com/tfr_mqc/report.html for the TfR RNAi samples. Dimensionality reduction was performed with the MDS algorithm implemented in Scipy after log2 transform of the read counts of the top 500 expressed gene. Differential Gene Expression analyses were carried out with edgeR using generalised linear models (GLM). The p-values of the test were corrected with the topTags function in R using the Benjamini–Hochberg method.

### RNA and protein stability assays

For mRNA stability assays, ∼5×10^6^ cells total were pre-treated with DFO for 5 hr and then incubated with Sinefungin (5 µg/ml, 5 min, 37°C) to inhibit trans-splicing and Actinomycin-D (10 µg/ml) to block transcription. Cells were harvested at various timepoints, RNA isolated, reverse transcribed, and quantified by qRT-PCR as described above. For protein stability assays, following DFO treatment, cells were incubated with cycloheximide (CHX, 100 µg/ml) to inhibit protein synthesis, harvested at various timepoints and analysed by western blots with anti-TfR, anti-HA or anti-BiP antibodies as loading control. Wherever used, cells were treated with FMK024 (FMK; 20 μM) or MG132 (MG; 25 μM) as indicated. Samples (1×10^7^) were processed for western blotting as described below. For TfR RNAi rescue experiments, cells were induced for RNAi for 24 or 48 hr, and then supplemented with holo-transferrin (Sigma Aldrich, 200 μg/ml), Lactoferrin (Sigma Aldrich, 200 μg/ml), or FeCl_3_ (F, 25 μM).

### Western blotting and antibodies

Total protein was extracted from 1×10^7^ cells in 1x SDS loading buffer, separated by SDS PAGE, transferred to PVDF membranes and immunoblotted with appropriate antibodies for 1 hr at room temperature: anti-BIP (rabbit, 1:5000), anti-TfR (rabbit, 1:5000), anti-Hsp70 (rabbit, 1:5000), anti-Ty (mouse, 1:2000) were generous gifts from Prof. J. D. Bangs (University at Buffalo, USA), and anti-HA (monoclonal HA-7, 1:5000, Sigma-Aldrich). The membranes were washed 3x in 1xPSBT and incubated for 1 hr at room temperature with the appropriate horseradish peroxidase (HRP)-conjugated secondary antibodies (anti-mouse IgG, 1:5,000; and anti-rabbit IgG, 1:5,000 dilution), developed after 1 min incubation with Super Signal ECL reagent (Thermo Fisher Scientific), and imaged using a ChemiDoc Gel Imaging MP System (Bio-Rad). Blots were quantified by densitometry using Image Lab software (Bio-Rad) as described in the appropriate figure legends. All statistical analyses were performed by one-way ANOVA or Student *t* test using PRISM v9 (GraphPad Software, Inc., San Diego CA). Differences were considered statistically significant at a *p-*value <0.05. The *n* values indicate the numbers of independent biological replicates performed and are shown in figure legends.

### Immunofluorescence microscopy and DAPI staining

Immunofluorescence assays were performed as previously described in (30). αHA (mouse) antibody dilution was 1:1000. Slides were washed with 1x PBS then incubated with secondary antibody for 1 hr at room-temperature: Alexa-488 conjugated α-mouse (1:500) (Molecular Probes, USA). Slides were washed in 1x PBS and mounted in VectaShield mounting medium (Vector Laboratories) containing 4′,6-diamidino-2-phenylindole (DAPI), sealed with coverslips and imaged on a Zeiss AxioImager M2 with an ORCA Flash 4 camera.

### Cell fractionation

HA::RBP5 cells were harvested, hypotonically lysed in distilled H_2_O plus protease inhibitor cocktail (PIC). Cytosolic and membrane fractions were separated by centrifugation (13,000g for 10 mins at 4 ^0^C). Total, cytosolic and organelle-associated fractions (10^7^ cells equivalents/fraction) were resuspended in 1xSDS sample buffer for standard SDS PAGE and immunoblotting with appropriate antibodies.

### Flow cytometry

∼5×10^6^ cells per condition were harvested at 4 °C, washed in 1x PBSG then resuspended in 100 μl 1x PBSG. Live cells were incubated with propidium iodide on ice for 10 minutes, washed in 1x PBSG and treated with 10 µg/ml RNaseA. The cells were incubated at 37 °C in the dark for 45 minutes, then analysed with an LSR Fortessa cell analyser using BD FACS Diva software (BD Biosciences). Endocytosis of Alexa488 conjugated Transferrin (Tf488) or FITC-conjugated Tomato lectin (TL-488) was carried out as previously described (30). Post-acquisition analysis of flow cytometry data was performed with FlowJo v10 software (BD Biosciences).

### Generation of inducible RNAi, RNAi-resistant reporter and overexpression cell lines

All TfR RNAi-sensitive and -resistant constructs were gifts from Prof. James D. Bangs (University at Buffalo). Generation of TfR RNAi cell lines and reconstitution of RNAi-resistant complements were performed as extensively described in our previous publications (29, 30). For inducible overexpression of wild type RBP5, the ORF 393 nts was amplified from genomic DNA (*T. brucei* strain 427) by PCR with flanking 5’-*Xho*l and 3’-*Nde*I restriction sites. The PCR amplicon was digested with *Xho*I/*Nde*I and cloned into the pLEW100v5x:Pex11 stem-loop vector (38). The resultant construct was confirmed by sequencing, linearised with *Not*I and transfected into a tetracycline inducible single-marker BSF cell line by electroporation as described in (68). Three clonal cell lines obtained by limiting dilution were selected for further analysis. To mutate Phe at position 61 to Ala, the gene was synthesised by GenScript Biotech (New Jersey, USA) changing nts at positions 181-183 from TTT to GCG with flanking *Xho*I/*Nde*I. The synthetic DNA was cloned into pLEW100v5x:Pex11 and transgenic cell lines generated as described above. The construct and cell line are referred to hereafter as F61A.

For inducible RBP5 RNAi, a 429 bp fragment (nts-50 to +379 relative to the RBP5 start codon) was PCR amplified with nested flanking 5’-HindIII/XbaI and 5’-XhoI/NdeI sites, sequentially digested first with NdeI/XbaI for antisense fragment and then with HindIII/XhoI for the sense fragment and cloned upstream and downstream of the Pex11 stuffer in the pLEW100v5x:Pex11 stem-loop vector (38) digested with the same enzymes. RNAi methodologies were performed as described above.

For Ty-tagged overexpression of RBP5, the coding region (nts 1-390) was amplified from genomic DNA (*T. brucei* strain 427) by PCR, using primers with 5’ Xhol and 3’ NdeI restriction sites, respectively. The 3’ primer introduced a Ty epitope tag sequence (EVHTNQDPLD) followed by TTA stop codon at the C terminus. Cloning into plew100.PEX11 vector were performed as described above and confirmed by sequencing.

### CRISPR/Cas9 epitope tagging and Knockouts

For knockouts and CRISPR/Cas9-mediated epitope-tagging of RBP5 at the N- and C-termini, linear DNA fragments were amplified from pPOT plasmids using oligonucleotides incorporating a 5’ overhang of ∼80 nucleotides of homology to the target gene and its adjacent UTR using primers and synthetic guide RNAs generated from LeishGEdit tool as described in (37). The amplicons were phenol-chloroform extracted and transfected into BSF cells constitutively expressing Cas9.

### Mutational sensitivity analyses of amino acid residues in RBP5

Using the RBP5 homology model derived from Phyre2, SuSPect and Missense3D were used to predict the phenotypic effects of missense mutations. Amino acid residues in RBP5 were ranked based on the mutation sensitivity score (9= high mutation sensitivity, 0= low) and amino acid variants were selected based on its predicted effect on function. Predicted solvent accessibility of each amino acid residue in wild type RBP5 generated by I-tasser was also considered in selection of variant amino acid.

### Data and software availability

All FASTQ files data are deposited at the NCBI SRA database under the bioproject accession number PRJNA671512. The code to reproduce the analysis pipeline and figures, the raw data and additional python scripts used for this study are available at GitHub https://github.com/mtinti/calvin_tfr and archived in Zenodo (https://zenodo.org/badge/latestdoi/285675864) for the DFO-treated samples and at https://github.com/mtinti/calvin_tfr2 and (https://zenodo.org/badge/latestdoi/285927639) for the TfR RNAi samples. The Jalview project file is available as supplementary material (def_analysis 2).

### RNA Immunoprecipitation (RIP)

was performed as described in Droll et al (69) Briefly, 5 x 10^8^ BSF cells were UV-cross-linked (400 mJ/cm^2^), flash frozen in liquid nitrogen and resuspended in 500 ul of lysis buffer (10 mM Tris pH 7.5; 10 mM NaCl; 0.1% NP-40) supplemented with 100 U RNasin, plus protease inhibitor cocktail (PIC). Lysis was achieved by passing through a 27G x 3/4 needle 20 times and pelleted at 6000 g. Twenty microlitres of the supernatant was used as input for Western blot analysis. The salt concentration was adjusted to 150 mM and incubated overnight at 4°C with anti-HA or anti-Ty agarose beads. After incubation, 20 uL and 100 uL of the flow through was reserved for Western blot and RNA analyses respectively. Antibody-bound beads were washed five times with ice-cold with IPP150, de-crosslinked with proteinase K (500 ul sample + 10 ul 10% SDS + 10 ul 0.4 M EDTA + 20 ul Proteinase K, 15 minutes at 42°C) and RNA was purified from flow-through and bound fractions using TriFAST. Eluted RNA (50 μ DNase-(SIGMA) and qRT PCR performed as described above. Relative fold-enrichment was determined by the equation 2^ΔCt^, where *C_t_* = *C_t_* (input) − *C_t_* (IP), C*_t_* = cycle threshold. The mock was either untagged parental cells or F61A mutant cells.

## Acknowledgements

This work was funded by a Wellcome Trust and Royal Society Sir Henry Dale Fellowship (208780/Z/17/Z) to CT. Initial financial support for RNA-Seq of TfR RNAi cells was generously provided by Prof. James D. Bangs at Department of Microbiology and Immunology, University at Buffalo (SUNY), USA. This provided critical preliminary data to support CT’s fellowship application. Prof. J. D Bangs also provided helpful advice, suggestions and various constructs and antibodies used throughout this manuscript.

Danielle C. Gordon is acknowledged for performing mutational analyses of RBP5 as part of her MRes project. We thank Drs. Martin Taylor (LSTHM), Nikolay Kolev, Cher-Peng Ooi, James Budzak, and Prof. Gloria Rudenko for helpful discussions and comments on the manuscript. We thank Prof. Sir Mike Ferguson (University of Dundee) for setting up collaborations with MT, who performed all Bioinformatic analyses. MT is supported by a Wellcome Trust Investigator Award (101842/Z13/Z) to Prof. Sir M. Ferguson. Prof. Mark Carrington (University of Cambridge, UK) is acknowledged for performing Northern Blot analyses and for helpful discussions.

---

**Figure S1. Generation of TfR RNAi cell line and validation of the iron starvation response** A tetracycline inducible RNAi cell line was generated targeting all subunits of native transferrin receptor E6 and E7. **a.** Growth curves of cells cultured in the presence (Tet+) or absence (Tet-) of 1 μg/ml tetracycline. Number of parasites/ml on the y-axis adjusted to starting density daily vs time on x-axis (days). Data are means ± SD (*n* = 3 technical replicates). **b.** TfR subunits were pulled down with holo-Tf conjugated beads from RNAi un-induced (tet-) or induced (tet +) cell lysates and detected by blotting with either anti-TfR (αTfR) at time points indicated. Anti-BiP (αBiP) serves as loading control. **c.** Relative TfR mRNA expression was analysed by qRT-PCR at indicated time points post RNAi inductions, normalized to un-induced cells at each time point. Data are means ± SD (*n* = 3 technical replicates, with 3 technical replicates for each *n*). **d.** Live cells were treated with tetracycline (Tet+) to initiate specific dsRNA synthesis or untreated (Tet-) and uptake of fluorescent-labelled holo-transferrin (Tf) or tomato lectin (TL) was analysed by flow cytometry. Normalised median fluorescent intensity (MFI) are shown for three independent biological replicates relative to un-induced controls. TL uptake serves as a surrogate control for general endocytosis. Data are means ± SD (*n* = 3 biological replicates). \**\*\*P* < 0.001, *\*\*P* < 0.002, as determined by Student t-test. **e.** qRT-PCR was performed to evaluate the effect of iron starvation following TfR RNAi on known homologues of iron-dependent proteins (IDPs). Relative mRNA levels of selected homologues of IDPs in BSF trypanosomes are shown. The relative quantity of each transcript was normalized against *ZFP3* (Tb927.3.720) relative to un-induced controls. **Abbreviations:** E6 (ESAG6), E7 (ESAG7), RBP5 (RNA-binding Protein 5; Tb927.11.12100), PAP2 (Phosphatidic Acid Phosphatase; Tb927.8.480), Hyp#1 (hypothetical protein; Tb927.8.490), Hyp#2 (hypothetical protein, Tb927.8.510); FeRed-1 (Ferric Reductase, Tb927.6.3320), FeRed-2 (Ferric Reductase, Tb927.11.4430); HpHbR (Haptoglobin Hemoglobin Receptor, Tb927.6.440), IRT1 (Iron Transporter, Tb927.11.8990); IRT2 (Iron Transporter, Tb927.11.9000); TAO (Trypanosome Alternative Oxidase, Tb927.10.7090), ACO (Aconitase), MLP (Mucolipin, Tb927.7.950), RNR1 (Ribonucleotide Reductase, Tb927.11.7840); RNR2 (Ribonucleotide Reductase, Tb927.11.12780). Data are means ± SD (*n* = 3 technical replicates, with 3 technical replicates for each *n*).

**Figure S2: Rescue of the iron starvation response by FeCl_3_ and genetic complementation a.** E6^R^ expressing cells were cultured in the presence of 1 μg/ml tetracycline (Tet +) supplemented with holo-transferrin (H, 200 μg/ml), Lactoferrin (L, 200 μg/ml), FeCl_3_ (F, 25 μM) or untreated (C) for 48 hr and total protein was extracted and blotted with αTfR or αBiP (loading control). **b.** E6^R^ expressing cells were pretreated with tetracycline for 24 hr followed by supplementation with FeCl_3_ for an additional 24 hr, and total protein blotted as in (a).

**Figure S3. Phylogenetic tree of Kinetoplastid RBP5 proteins.** Proteins with homology to RBP5 were retrieved from TritrypDB and the output was saved as public search: https://tritrypdb.org/tritrypdb/app/workspace/strategies/import/3ef3dfbc6f322bf6. The sequences were imported in Jalview and aligned with the Mafft alghoritm (L-INS-i) using the Jalview interface. An aligned region of 102 amino acids with no gaps was used to create a phylogenetic tree with the Jalview interface (Average Distance and BLOSUM62 matrix). The tree was exported in newick format and loaded in FigTree. Proteins with accession numbers: TM35_000312320 and TM35_000312310 (*Trypanosoma theileri*), PCON_0057560 (*Paratrypanosoma confusum*) and DQ04_03381000 (*Trypanosoma grayi*) cluster together but very divergent relative to the other putative RBP5 proteins. For this reason, the alignment was rerouted using these divergent sequences as outgroup.

**Figure S4 PCR validation of RBP5 C- and N-terminal epitope tagging a.** *In situ* N-terminal tagging of RBP5 with 6xHA retaining its native 3’-UTR. Selection of transgenic cells was achieved using puromycin (PAC) and Hygromycin (Hyg). Arrows indicate position of PCR primers to verify RBP5 locus-specific integration. The corresponding size fragments are shown by genomic DNA (gDNA) PCR from wild type (WT), single allele (Hyg) or both alleles (Hyg/Pac) tagged cell lines. All diagrams are not drawn to scale and *in situ* tags were done by CRISPR/Cas9-mediated PCR-based method published in (37). **b.** Schematic shows strategy for generating *in situ* C-terminally tagged RBP5 (RBP5::mNG-Ty) fused with *PFR* 3’-UTR and hygromycin (HYG) as drug selection cassette. Arrows indicate position of PCR primers to analyse correct integration with corresponding size fragments shown in agarose gel on the right. CRISPR/Cas9 mediated tagging strategy in (a) was used. **c.** Effect of iron starvation on C terminal epitope tagged RBP5 protein expression. Schematic of C-terminal *in situ* tagged RBP5 (RBP5::mNG-Ty) fused with PFR 3’-UTR. RBP5::mNG-Ty expressing cells were incubated with deferoxamine [25 μM, 5 hr or 5 μM, 24 hr]. Total protein was extracted and blotted with αTy or αBiP antibodies. RM, M indicate RBP5::mNG-Ty and mNG::Ty polypeptides, respectively. Bar chart (right) shows quantification of RBP5::mNG-Ty relative to BiP signals. **d.** Effect of iron starvation on *RBP5* mRNA abundance by Northern blot analyses Log phase BSF cells (∼5×10^5^ cells/ml) were incubated with deferoxamine (DFO), Iron (III) chloride (Fe) or untreated (Ctrl), RNA was isolated and analysed by Northern blotting with probes to full length *RBP5* ORF plus 1,235 bp downstream of the STOP codon. The blot was re-probed with 5.8S rRNA as loading control (70). For each sample, RBP5 mRNA levels were quantified relative to 5.8s rRNA and shown as Qty beneath the blot. In contrast to curated and experimental data of poly A sites from TritrypDB, these data (from a single experiment) show that the RBP5 3’-UTR is ∼1.291 Kb from the predicted STOP codon. Also note that RBP5 appears on the blot as single homogenous species suggesting no additional processing occurs following DFO treatment.

**Figure S5. *RBP5* RNAi and knockout Effect of *RBP5* RNAi on cell growth**. *RBP5* RNAi cells were counted using a haemocytometer for 72 hrs with (RBP5 RNAi) or without (Ctrl) 1 μg/ml tetracycline. The value on the y-axis represents measured value times the dilution factor. Data are means ± SD, *n* = 3. **a.** qRT-PCR analyses showing levels of *RBP5* mRNA with or without RBP5 RNAi at 48 hr, 1 μg/ml tetracycline. Data are means ± SD, *n* = 3 technical replicates. qRT-PCR analyses showing levels of *RBP5* and *TfR* mRNA following treatment with 1 μg/ml tetracycline for 25 hr (RNAi), followed by 25 μM DFO for 5 hr (DFO), combined DFO and tetracycline (DFO/RNAi) or no treatment (Ctrl). Data means ± SD, *n* = 5 biological replicates, with 3 technical replicates for each *n*. **b.** Effect of *RBP5* RNAi on HA::RBP5 protein levels. *RBP5* RNAi construct was transfected into HA::RBP5 expressing cells, RNAi was induced for 24 hr (RNAi) or not (Ctrl) and protein levels determined by western blots with either anti-HA (R, RBP5) or anti-BiP (B, loading control) antibodies. **c.** Generation and PCR validation of *RBP5* knockout cell lines. The schematic shows the targeting strategy of the *RBP5* wild type locus. The arrows indicate the position of oligonucleotide primers used for PCR confirmation of the inserted drug resistance cassettes: Hygromycin (HYG) and Puromycin (PAC). The agarose gel images on the right show PCR amplicons obtained using genomic DNA template derived from either wild type (Par), *RBP5* single knockout with HYG (sKO), or *RBP5* double knockout (RBP5-/-) with HYG and PAC cassettes. The colours indicate the primers used as shown on the schematic. Four *RBP5-/-* clonal cell lines were tested from two independent transfections. In all instances, integration of both HYG and PAC cassettes were confirmed by PCR. However, the *RBP5* ORF had been duplicated in the genome as confirmed by a PCR product using primers that anneal to the ORF.

**Figure S6. Multiple Sequence Alignment of RBP5 RRM domain a.** Selected RRM-1 Family (PF0076) proteins from different species were aligned with the RRM domain of *T. brucei* RBP5. Arrowhead shows conserved Phe residue RNP-1 motif. The residue was mutated to Ala based on conservation analysis using Jensen-Shannon divergence, structural and mutation analyses by SuSPect and Missense 3D. Mutation to alanine maintains hydrophobicity at this position potentially disrupting binding to target RNA sequence (56). Abbreviations: PABP: poly(A) binding protein; SCHOP: *Schizosaccharomyces pombe*: DROME: *Drosophila melanogaster*; ELAV: Embryonic Lethal Abnormal Vision; TIA: T-cell-restricted intracellular antigen-1; U2AF: general splicing factor; U2AF2: U2 Small Nuclear RNA Auxiliary Factor 2; RU2B: small nuclear ribonucleoprotein polypeptide B2; SNRPA: U1 small nuclear ribonucleoprotein A; PRP24: Precursor RNA processing, gene 24. **b.** Radial tree of multiple sequence alignment in (a). The tree was generated using CLC Genomics Workbench 20 with default parameters.

**Figure S7. *RBP5* mRNA enrichment by RIP qRT-PCR a.** BSF cells expressing c-terminally Ty-tagged wild type RBP5 (RBP5^WT^), an RRM mutant (RBP5^F61A^) or untagged Parental (Par) strain were treated with tetracycline, supplemented with deferoxamine (DFO, bottom) or without (Control, top) for 4 hours. RIP was performed with anti-Ty beads, followed by immunoblotting with anti-Ty (αTy, R) and anti-BiP (αBiP, B). **b.** Bar chart shows relative fold-enrichment of RBP5 mRNA levels quantified by qRT-PCR and corrected for background either with untagged Par (RBP5^WT^ vs Par) or with RBP5^F61A^ (RBP5^WT^ vs RBP5^F61A^). The value shown on the y-axis is normalised to 28s rRNA as non-bound control. Note that: (i) there was no enrichment of *RBP5* mRNA in both untagged Par and RBP5^F61A^ in either condition, as expected, (ii) bearing in mind that RBP5 protein is overexpressed, we see a 20 – 40-fold RBP5 mRNA enrichment in Ctrl cells (blue bars). The data suggest that increased RBP5 protein expression is sufficient to elicit RBP5 protein/mRNA association. No conclusion can be made whether iron depletion by DFO in the context of overexpression enhances the interaction between RBP5 mRNA and protein since these data are derived from a single experiment. However, the data in Fig 8 showing endogenous HA-tagged protein levels reflects a physiological scenario.

