## Supplementary figures and images for "Novel aspects of iron homeostasis in pathogenic bloodstream form *Trypanosoma brucei*"

### Fig S1

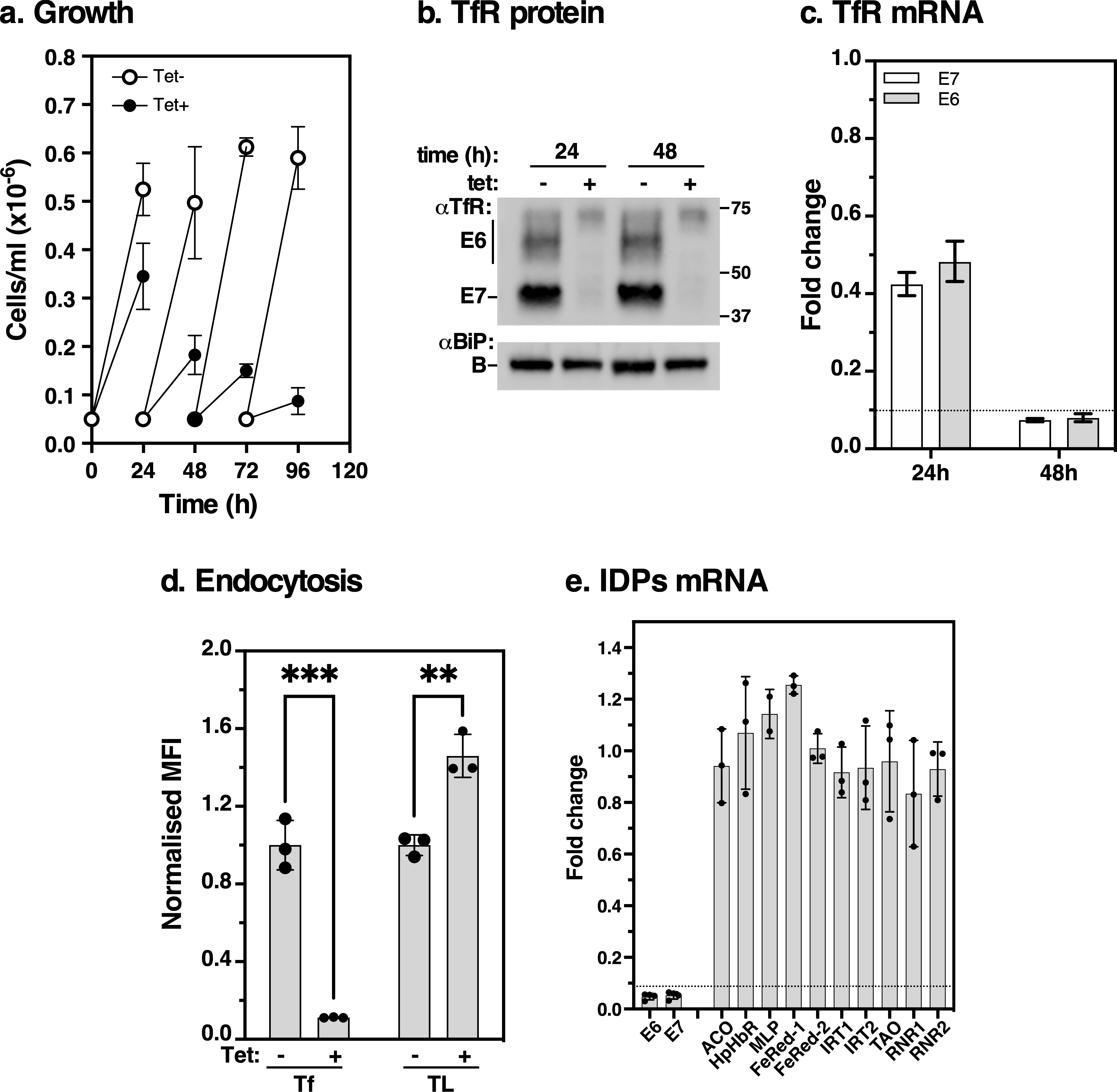

### Fig S2

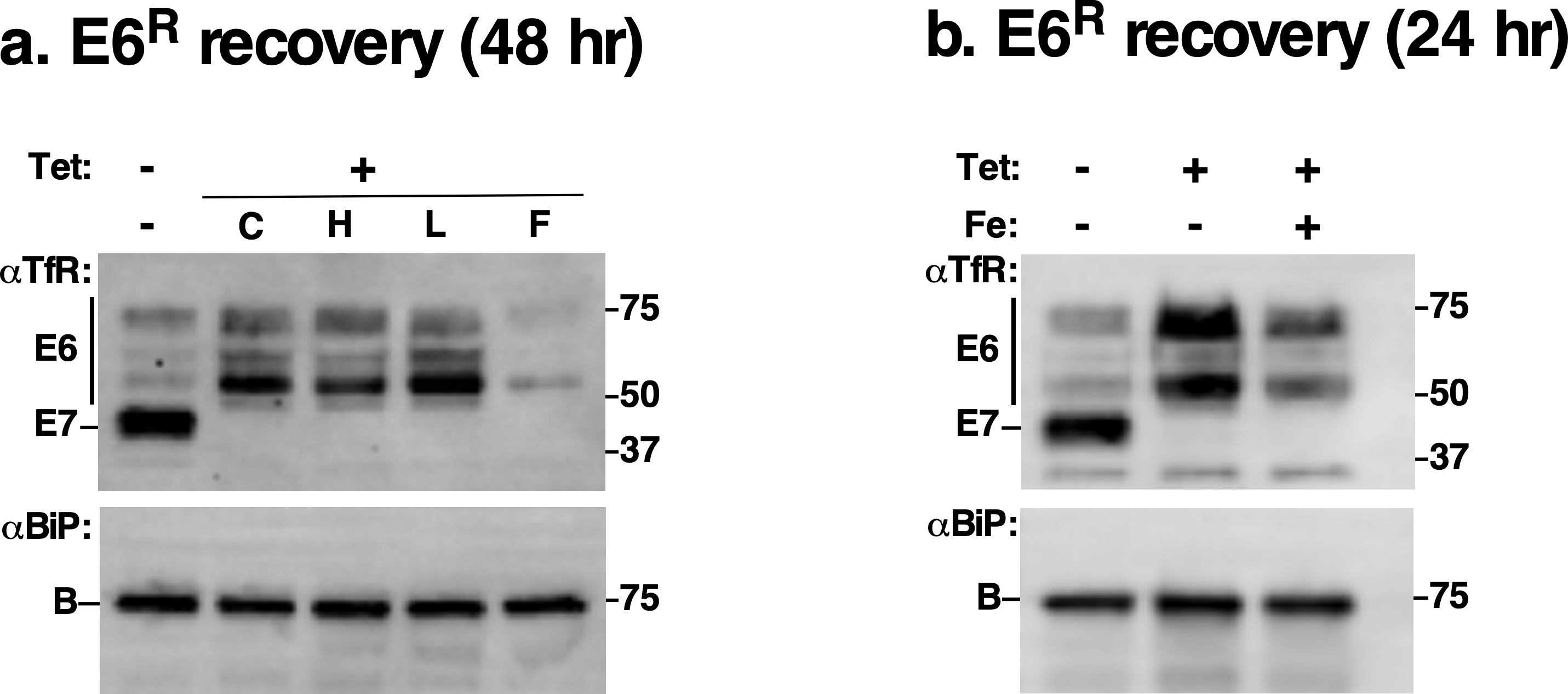

### Fig S3

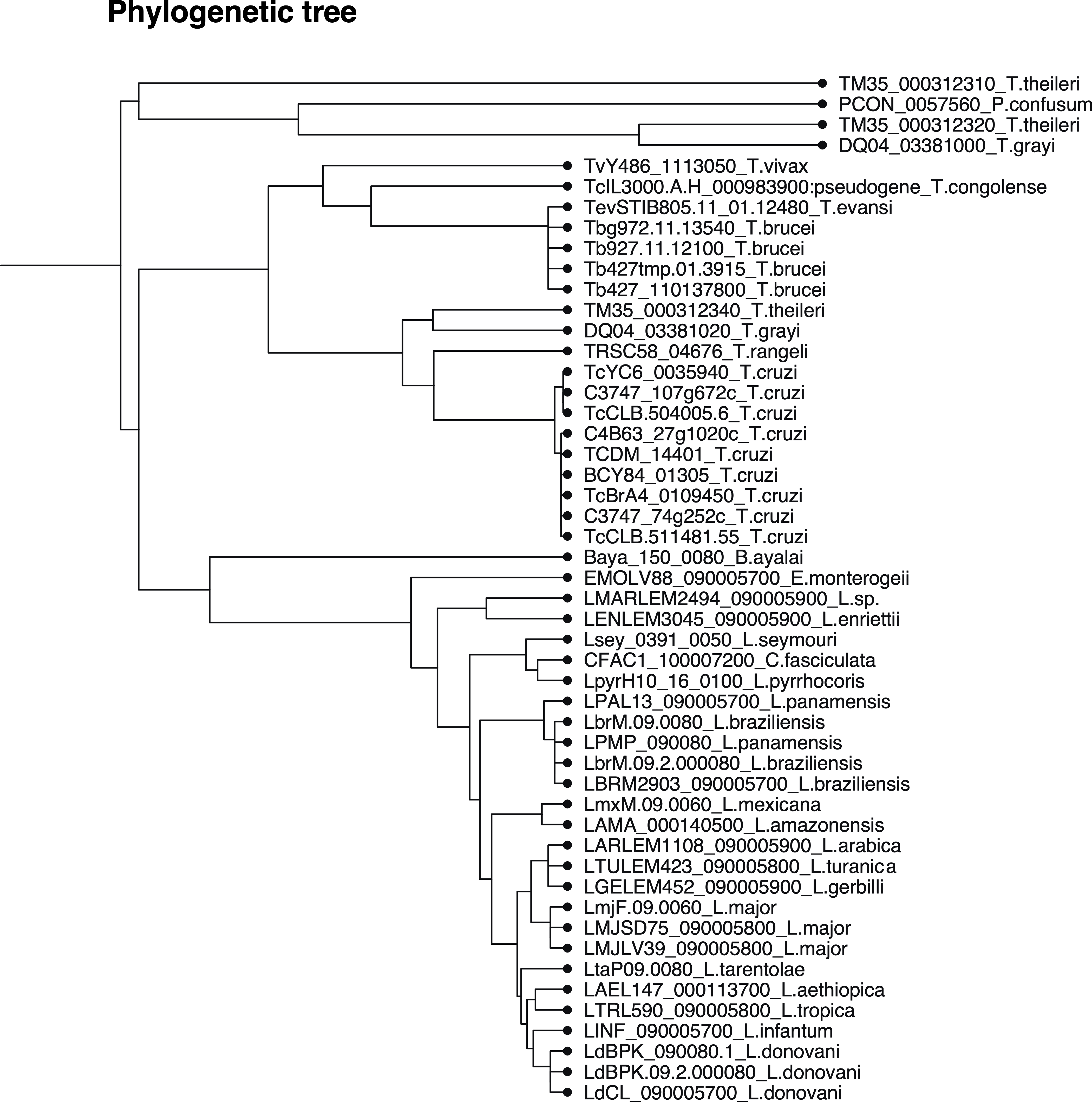

### Fig S4

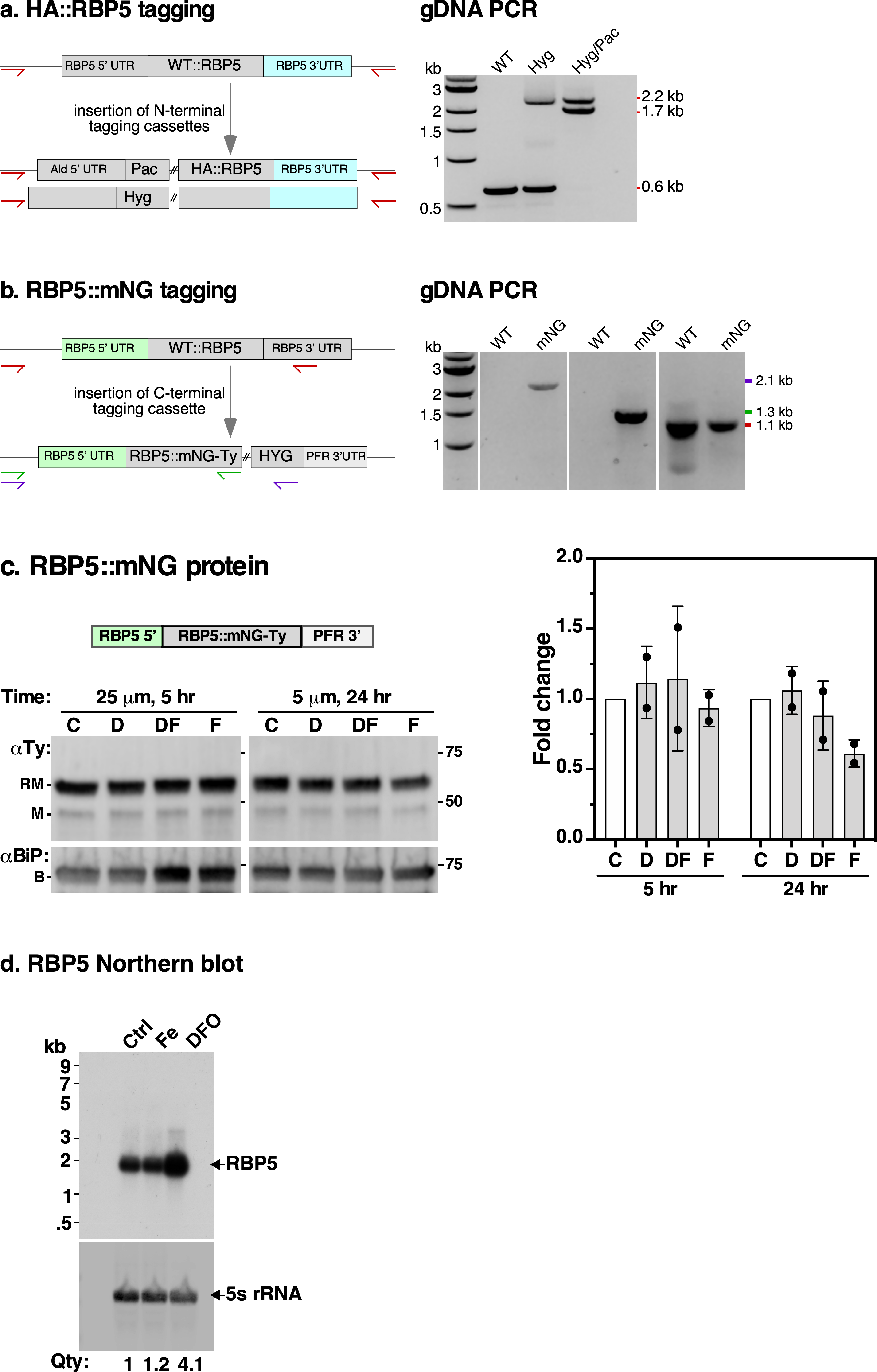

### Fig S5

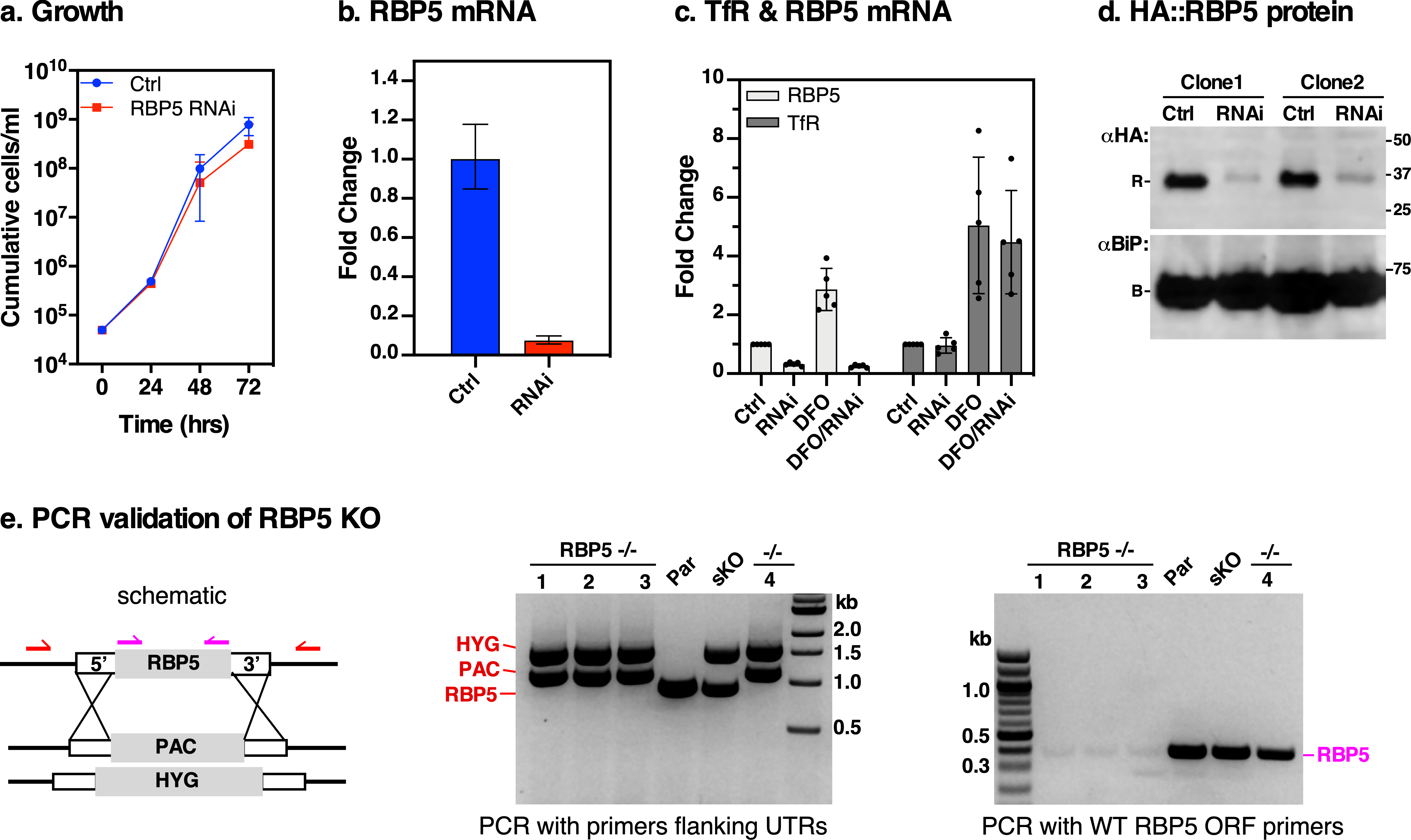

### Fig S6

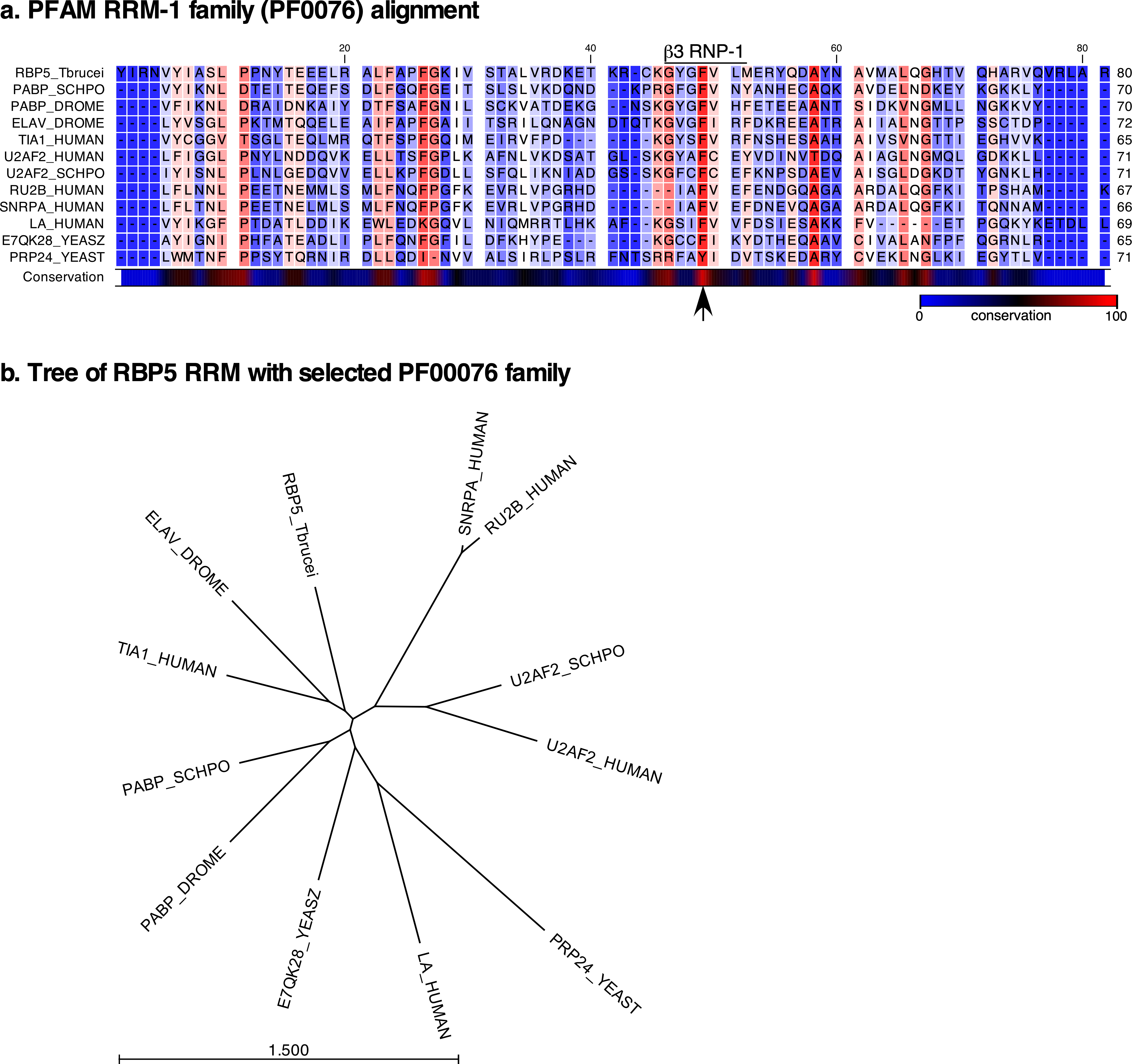

### Fig S7

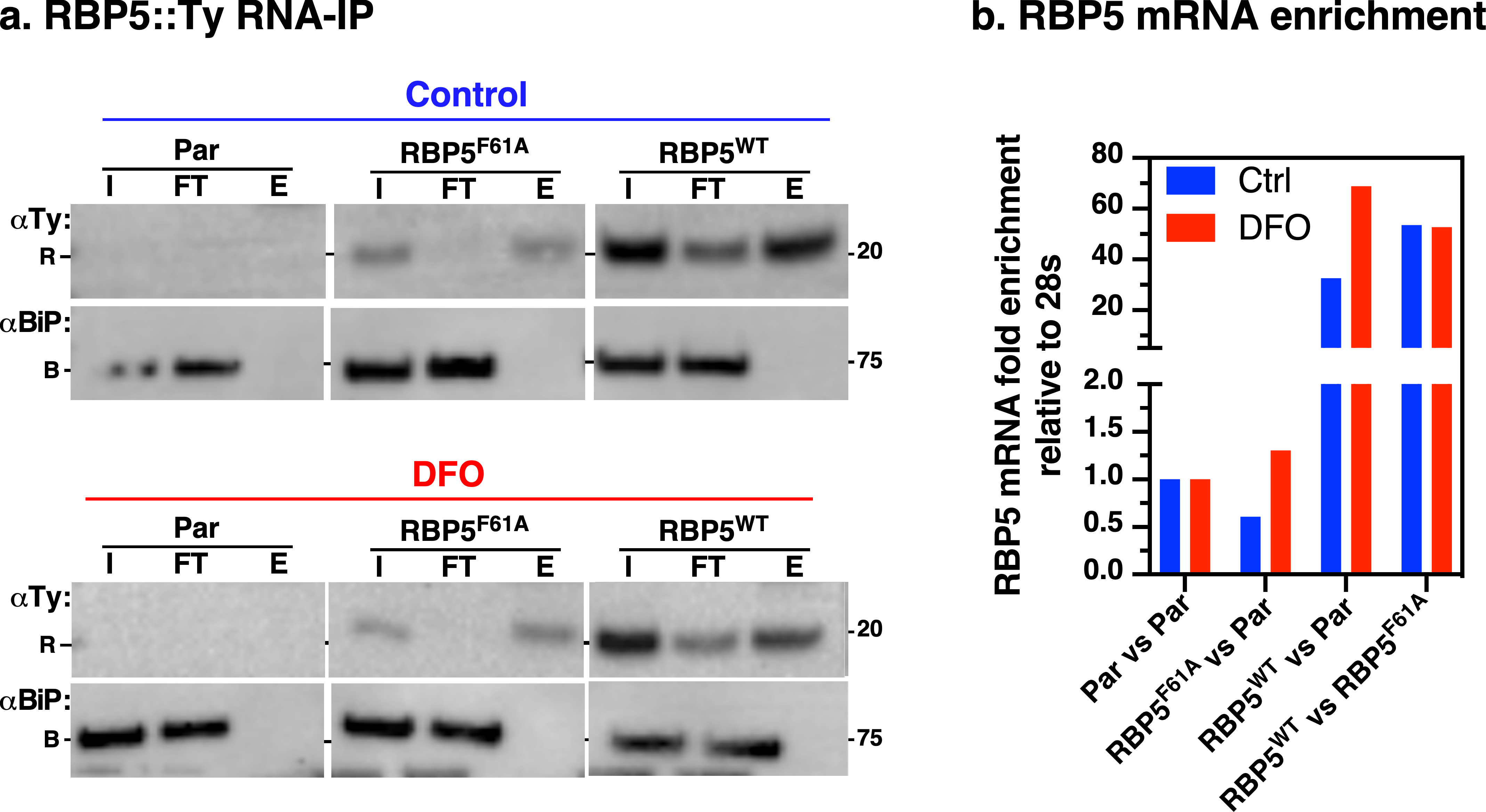
